# Modelling quantitative fungicide resistance and breakdown of resistant cultivars: designing integrated disease management strategies for Septoria of winter wheat

**DOI:** 10.1101/2022.08.10.503500

**Authors:** Nick P Taylor, Nik J Cunniffe

## Abstract

Plant pathogens respond to selection pressures exerted by disease management strategies. This can lead to fungicide resistance and/or the breakdown of disease-resistant cultivars, each of which significantly threaten food security. Both fungicide resistance and cultivar breakdown can be characterised as qualitative or quantitative. Qualitative (monogenic) resistance/breakdown involves a step change in the characteristics of the pathogen population with respect to disease control, often caused by a single genetic change. Quantitative (polygenic) resistance/breakdown instead involves multiple genetic changes, each causing a smaller shift in pathogen characteristics, leading to a gradual alteration in the effectiveness of disease control over time. Although resistance/breakdown to many fungicides/cultivars currently in use is quantitative, the overwhelming majority of modelling studies focus on the much simpler case of qualitative resistance. Further, those very few models of quantitative resistance/breakdown which do exist are not fitted to field data. Here we present a model of quantitative resistance/breakdown applied to *Zymoseptoria tritici*, which causes Septoria leaf blotch, the most prevalent disease of wheat worldwide. Our model is fitted to data from field trials in the UK and Denmark. For fungicide resistance, we show that the optimal disease management strategy depends on the timescale of interest. Greater numbers of fungicide applications per year lead to greater selection for resistant strains, although over short timescales this can be offset by the increased control offered by more sprays. However, over longer timescales higher yields are attained using fewer fungicide applications per year. Deployment of disease-resistant cultivars is not only a valuable disease management strategy, but also offers the secondary benefit of protecting fungicide effectiveness by delaying the development of fungicide resistance. However, disease-resistant cultivars themselves erode over time. We show how an integrated disease management strategy with frequent replacement of disease-resistant cultivars can give a large improvement in fungicide durability and yields.

**Author Summary:** Plant pathogens pose a major threat to crop yields. The two most common forms of pathogen control, namely use of fungicides and deployment of disease resistant cultivars, are threatened by pathogen evolution causing fungicide resistance or erosion/breakdown of cultivar control. There are two categories of resistance/breakdown; qualitative or quantitative. Although resistance to many cultivars and the most common fungicides is quantitative, the mathematical modelling literature focuses almost exclusively on qualitative resistance, for simplicity or due to lack of appropriate data required to fit a model of quantitative resistance. In this study we present the first model focusing on both quantitative fungicide resistance and cultivar breakdown to be fitted to field data. We use the disease of wheat, Septoria leaf blotch, as a case study. After fitting our model to field trial data from the UK and Denmark, we use it to demonstrate how to design sustainable disease management strategies that optimise yield. We show that combining resistant cultivars with fungicide applications can prolong the effectiveness of both strategies, but that the optimal number of fungicide applications depends on the timescale of interest. Over short timescales, the optimal strategy uses more fungicide applications per year than over longer timescales.

## Introduction

Plant pathogens are a significant threat to global food security (***Ristaino et al., 2021***; ***Savary et al., 2012***). Diseases caused by plant pathogens routinely lead to large losses in crop yields (***Savary et al., 2019***), and significant proportion of growers’ time and money can therefore be spent on disease management (***Schumann and D’Arcy, 2010***). Spraying crops with fungicide and using cultivars which are genetically resistant to pathogens are two of the most effective control strategies. However, each presents huge selection pressure on pathogen populations, which consequently evolve in response (***Zhan et al., 2015***). This leads to pathogen populations which exhibit fungicide resistance (***Brent and Hollomon, 2007***), as well as the loss of effectiveness (‘breakdown’) of disease resistant host cultivars (***McDonald and Linde, 2002***). Mathematical models have an important role in understanding the processes underlying these changes (***Cunniffe et al., 2015***), which involve complex interactions occurring over long time scales.

Both fungicide resistance and breakdown of disease resistant crop cultivars can be characterised as qualitative or quantitative (***McGrath, 2007***; ***Didelot et al., 2016***). Qualitative resistance/breakdown is usually caused by a single genetic change leading to a major shift in the characteristics of the pathogen population of interest (***Lucas et al., 2015***). Qualitative resistance/breakdown is therefore sometimes called monogenic (***St. Clair, 2010***). For some fungicides, a single genetic change in the pathogen leads to a step change in the fungicide sensitivity of affected individuals. For resistant cultivars, qualitative breakdown tends to occur in cases in which disease resistance depends on a single resistance gene. A single pathogen mutation can then lead to the cultivar suddenly becoming susceptible to disease. For both fungicides and host plant resistance, qualitative resistance/breakdown involves a very sudden transition from good to bad control, even if the control had previously been effective for several years.

In contrast, quantitative resistance/breakdown is characterised by small, gradual shifts in the make-up of the pathogen population, with multiple genetic changes which accumulate over time (***McGrath, 2007***; ***Didelot et al., 2016***). For fungicides often these changes are polygenic, although in some cases multiple mutations in the same gene can lead to a gradual loss of effectiveness in control. Reduced susceptibility of modern disease resistant cultivars is often achieved by stacking or pyramiding minor resistance genes (i.e. combining more than one gene for resistance), each with a small effect on disease susceptibility. This means that the pathogen population gradually attains resistance, and since mutations in different avirulence genes are involved, this involves polygenic changes in the pathogen population. In both cases, the typical pattern is control that has partial, but diminishing, effect for several growing seasons until eventually it becomes too ineffective for adequate control in the field.

Quinone-outside inhibitor fungicides (QoIs) provide an example of qualitative (monogenic) fungicide resistance. In particular, the G143A mutation in *cytb* (cytochrome b gene) is linked to high levels of resistance to QoI fungicides that disrupt electron transport during cellular respiration (***Estep et al., 2013***). Taking an example from the important wheat disease Septoria tritici blotch (STB, caused by *Zymoseptoria tritici*), QoIs were very effective when they were first introduced in the 1990s. However, after only a few years of use they became ineffective (***Blake et al., 2018***), and the shift from good to poor control happened very rapidly. In 2002, approximately 80% of fungicide sprays contained QoIs in a wheat disease monitoring program in northern Germany (***Beyer et al., 2011***). After breakdown of QoI efficacy, their use dropped to around 4% in 2006. Contemporary fungicide spray programmes targeting control of STB tend to rely upon azole and succinate dehydrogenase inhibitor (SDHI) fungicides (***Kirikyali et al., 2017***; ***Torriani et al., 2015***), to which resistance is quantitative. For azoles, multiple mutations in the CYP51 enzyme are most often implicated (***Cools and Fraaije, 2013***; ***Kristoffersen et al., 2020a***).

Often growers combine fungicides and resistant cultivars, sometimes as part of so-called ‘integrated disease management’ strategies. The philosophy, closely linked to ‘integrated pest management’ (IPM), emphasises simultaneous deployment of multiple control strategies. The underpinning idea is to obtain more effective control but with less dependence on any single strategy, particularly chemicals (***Barzman et al., 2015***). In agriculture in the developed world, growers routinely combine resistant cultivars with fungicide spray programmes. Clearly this has the potential to lead to more effective control in any single season. However, the theory of fungicide resistance indicates any factor causing an overall reduction in pathogen growth rate of both resistant and sensitive pathogen strains leads to slower selection and so an increased lifetime of the chemical of interest (***van den Bosch et al., 2014***). Recent theoretical work has applied this to show how fungicide sprays can also make host resistance more durable (***Carolan et al., 2017***).

In general, mathematical models have played a central role in understanding how control via fungicides and cultivar resistance can be optimised (***Cunniffe et al., 2015***; ***Corkley et al., 2022***). Until now, mathematical modelling has primarily focused on qualitative resistance/breakdown (***Hobbelen et al., 2011a***,b, ***2013***; ***Mikaberidze et al., 2017***; ***Elderfield et al., 2018***; ***Fabre et al., 2015***; ***Watkinson-Powell et al., 2020***; ***van den Bosch and Gilligan, 2003***; ***Fabre et al., 2012***), with very few focusing on quantitative resistance (***Shaw, 1989***; ***Miele et al., 2021***; ***Fabre et al., 2022***). However, resistance to many fungicides and resistant cultivars currently in use is quantitative. Despite the benefits of integrated disease management, few models in the literature address both fungicide and cultivar resistance, with most focusing on either fungicide (***van den Berg et al., 2013***; ***Hobbelen et al., 2011b, 2013***; ***Elderfield et al., 2018***) or cultivar (***Mikaberidze and McDonald, 2020***; ***Fabre et al., 2015***; ***Delmas et al., 2016***; ***Fabre et al., 2022***) control. In this paper we present a model of quantitative resistance incorporating both fungicide and disease-resistant cultivar control.

In the modelling literature for fungicide resistance, evolution is typically split into an ‘emergence phase’ and a ‘selection phase’. The emergence phase is the period during which resistant strains invade the pathogen population following a mutation event (***van den Bosch et al., 2011***). The selection phase is the period after the emergence phase, in which strains with different fungicide sensitivities are present in the population, and when resistant strains are selected for by fungicide application. The models currently in the literature address either the emergence phase (***Mikaberidze et al., 2017***) or the selection phase (***Hobbelen et al., 2013***; ***Elderfield et al., 2018***). The model presented in this paper instead represents both phases of resistance development in a single model framework.

Plant disease systems are inherently stochastic, most obviously because of the effects of weather on pathogens’ spread (***Agrios, 2004***). However, few models of fungicide resistance or cultivar susceptibility incorporate environmental stochasticity, despite some exceptions (***te Beest et al., 2013***; ***Willocquet et al., 2020***; ***Lo Iacono et al., 2013***). We aim to understand the impact of environmental stochasticity on the optimal disease management recommendation. Using environmental variability in the model is not only inherently more realistic, but also helps us understand the best- and worst-case scenarios for proposed disease management strategies.

To the best of our knowledge, the model we present here is the first that focuses on quantitative resistance and is fitted to field data for both cultivar and fungicide control. We use STB, the most prevalent disease of wheat worldwide (***Torriani et al., 2015***; ***Suffert et al., 2011***), as our case study. It is a particularly economically important disease; the fungicide market in Europe alone is estimated to be more than $2.4bn (***Torriani et al., 2015***), of which $1.2bn is primarily targeted towards managing STB. Assessments of control efficacy against STB are routinely taken during field trials of new cultivars (***Kristoffersen et al., 2020b***), and in fungicide resistance assessments (***Blake et al., 2018***). Crucially these programs span multiple years, providing us with a source of the type of data we require to fit our model.

In this paper we use a model of quantitative fungicide resistance and breakdown of resistant cultivars to explore the optimal way to combine host and fungicide control strategies to minimise disease impact and maximise yield. The model includes pathogen mutation, initial standing variation in the pathogen population and environmental stochasticity, and is the first model of quantitative resistance in the literature that is fitted to field data. We use our model to address the following questions:

1. How does the number of fungicide applications per year affect resistance development, disease severity and yield over time?
2. How does environmental stochasticity influence the optimal strategy?
3. How does the optimal strategy depend on the time-frame of interest?
4. How can resistant cultivars be optimally combined with fungicide control for an effected integrated disease management approach?

## Materials and methods

### Model structure

Our model tracks the disease caused by, and evolution of, a pathogen population over multiple growing seasons. Within a single growing season, we model the growth of the pathogen population and the loss of healthy tissue. ‘Disease severity’ is infected tissue measured as a percentage of the maximum leaf area (with the other tissue assumed to be healthy, and measured in the same way). For simplicity, there is no consideration of spatial effects.

The model is an extension of a simple Susceptible-Infected disease model (*SI*) which includes a more detailed description of the pathogen population. This allows us to explore the evolutionary dynamics of the pathogen in greater detail. In particular, we consider the level of resistance to the fungicide as a continuous spectrum. This allows us to explore the effect of using new fungicides in the face of quantitative pathogen evolution and/or disease-resistant cultivars. Initially we focus on fungicide control only, before exploring the effect of both fungicide and cultivar control. A full description of model variables is found in Table 1.

**Table 1.** List of variables used. The variables *S, I* and the integral 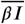 all vary with time *t*. Infection *I* also depends on trait values *k, l*. The pathogen infection rate *β* depends on *k, l* and *t*. The uncontrolled pathogen infection rate *β*_0_ depends on the disease pressure in a particular year.

| Variable (fungicide only model) | Variable (full model) | Description |
| --- | --- | --- |
| $S(t)$ | $S(t)$ | Susceptible host tissue |
| $I(k, t)$ | $I(k, l, t)$ | Infected host tissue |
| $P(a, b)$ | $P(a, b)$ | Probability of mutation from parent $b$ to offspring $a$ |
| $G(k, t)$ | $G(k, l, t)$ | Per capita growth rate |
| $t$ | $t$ | Time |
| $k$ | $k$ | Fungicide sensitivity trait |
| $-$ | $l$ | Host susceptibility trait |
| $\beta_0$ | $\beta_0$ | Pathogen infection rate (no control) |
| $\beta(k, t)$ | $\beta(k, l, t)$ | Pathogen infection rate (control applied) |
| $g(t)$ | $g(t)$ | Host growth function |
| $\Gamma(t)$ | $\Gamma(t)$ | Host senescence function |

#### Single strain model (fungicide only)

We first consider a single pathogen strain, for the purposes of explanation, before introducing the full model. We assign this pathogen strain a ‘fungicide trait value’ *k*, which describes its fitness with respect to the fungicide. The trait value *k* is between 0 and 1, with 0 being fully sensitive (i.e. the pathogen is fully controlled by a fungicide application, at least directly after it is sprayed) and 1 being fully resistant (i.e. completely unaffected by a fungicide application). We use ‘strain *k*’ as a shorthand for ‘the pathogen strain with trait value *k*’.

The density of susceptible host at time *t* is denoted by *S*(*t*), and the density of strain *k* at time *t* is denoted by *I*(*k, t*). The rate of loss of healthy (susceptible) leaf tissue is given by:

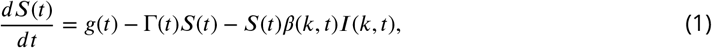

where *g*(*t*), Γ(*t*) and *β*(*k, t*) are host growth, host senescence and the pathogen infection rate respectively, which are all described below.

The rate of increase of disease severity (of strain *k*) is given by:

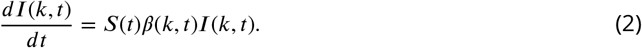

The pathogen infection rate, *β*(*k, t*), depends on the pathogen response to the fungicide. ***Hobbelen et al. (2013***); ***Elderfield et al. (2018***); ***Taylor and Cunniffe*** (***2022***) use a fungicide dose response of the form:

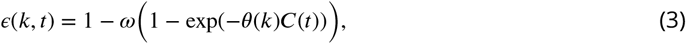

where *ϵ* is the proportionate effect of the fungicide on strain *k, ω* is the asymptote of the dose response curve and *θ*(*k*) is the curvature of the dose response curve with respect to the pathogen, which here also depends on the strain *k*. We choose the asymptote parameter, *ω*, to equal 1, as was used for high risk fungicide in ***Elderfield et al. (2018***). This means that the infection rate at theoretical infinite dose is 0.

The relationship between *k* and *θ*(*k*) is that *θ*(*k*) = − log(*k*). Then, since we have assumed the asymptote *ω* = 1, we find:

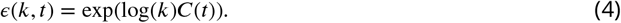

This means that the proportionate effect of the fungicide on the infection rate of this pathogen strain is *k* when the concentration is 1. In general the effect depends on the fungicide concentration, which is assumed to decay exponentially with time *t*, at rate Λ, after each application as in ***Elderfield et al. (2018***); ***Hobbelen et al. (2013***). The fungicide concentration equation is:

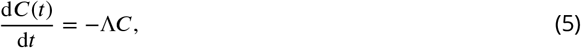

where *C* is initially 0 but instantaneously increases by 1 every time the fungicide is applied. We choose a decay rate of Λ = 0.5 ×(6.91 + 11.1) × 10^−3^ = 9.005 × 10^−3^ degree-days^-1^, which is the average of the two fungicide decay rates in ***Elderfield et al. (2018***).

The fungicide effect acts as a multiplier on the underpinning infection rate *β*_0_. When the concentration is 0, the effect is 1, so when there is no fungicide present all strains behave identically. When the fungicide is applied the infection rates of strains with lower *k* values are suppressed more than that of strains with higher *k* values:

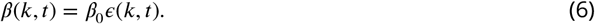

The fungicide can be applied at up to three time points: *T*_1_ (growth stage 32), *T*_2_ (growth stage 39) and *T*_3_ (growth stage 61) (***van den Berg et al., 2013***; ***Elderfield et al., 2018***). A one treatment programme is applied at *T*_2_, a two treatment programme is applied at *T*_2_ and *T*_3_, a three treatment programme is applied at *T*_1_, *T*_2_ and *T*_3_ (Table 2).

**Table 2.** Times that are required in running the model. Here we use Zadoks’ growth scale (***Zadoks et al., 1974***). The times *T*_1_, *T*_2_ and *T*_3_ are as in ***van den Berg et al. (2013***); ***Elderfield et al. (2018***). The start time is growth stage 32, corresponding to the emergence of leaf 3 (*T*_1_). The final time *t*_*end*_ (growth stage 75) is interpolated from the value at growth stage 61 (*T*_*GS*61_ = *T*_3_ = 2066 degree-days) and growth stage 87 (*T*_*GS*87_ = 2900 degree-days) from ***van den Berg et al. (2013***); ***Elderfield et al. (2018***). This assumes that the temperature remains approximately constant in the late stages of the season. We also round the values to the nearest integer. Growth stage 75 was chosen for the end time because it was the time in Dataset D which links yield to disease severity (Table 3). Any host or pathogen growth before *t*_*start*_ can be scaled into the amount of inoculum *I*_0_ in the model at *t*_*start*_; the procedure used to fit the initial inoculum parameter, *I*_0_, is reported in Appendix 1.

| Parameter | Growth stage | Time (degree-days) | Purpose |
| --- | --- | --- | --- |
| $t_{start} = T_1$ | 32 | 1456 | Application programme: 3; also start time |
| $T_2$ | 39 | 1700 | Application programmes: 1, 2, 3 |
| $T_3$ | 61 | 2066 | Application programmes: 2, 3 |
| $t_{end}$ | 75 | 2515 | End time |
| $T_{GS61}$ | 61 | 2066 | Start of senescence in *Elderfield et al. (2018)* |
| $T_{GS87}$ | 87 | 2900 | End time in *Elderfield et al. (2018)* |
| $T_1$ | 32 | 1456 | Fitting $I_0$ |
| $t_A$ | 63 | 2130 | Fitting $I_0$ |
| $t_B$ | 71 | 2387 | Fitting $I_0$ |
| $t_{end}$ | 75 | 2515 | Fitting $I_0$ |

The host is assumed to grow within the season before a period of senescence. We use the same form for growth and senescence as in ***Hobbelen et al. (2011b***); ***Elderfield et al. (2018***). The growth equation is:

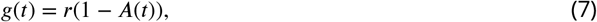

where *A*(*t*) is the total amount of healthy and infected tissue. The value for *r* is 0.0126 degree-days^-1^ as in ***Hobbelen et al. (2011a***); ***Elderfield et al. (2018***). We non-dimensionalised so that the maximum leaf area is 1, as in ***Taylor and Cunniffe*** (***2022***). The initial total quantity of host tissue is 0.05*/*4.1, using the values for initial host area and maximum host area from ***Hobbelen et al. (2011a***).

The senescence equation, which controls the time-dependent rate at which leaf senescence occurs, is:

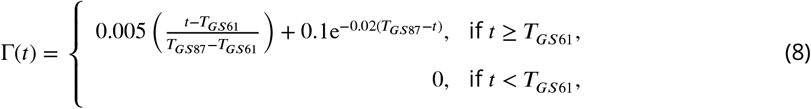

as in ***Hobbelen et al. (2011a***); ***Elderfield et al. (2018***) – see Table 2.

#### Model accounting for diversity within the pathogen population (fungicide only)

The population model, which accounts for diversity within the pathogen population (Table 1) addresses a continuous spectrum of pathogen strains with trait values (*k*) varying from 0 to 1. It generalises the single strain model to an entire population of different pathogen strains with differing initial densities. Healthy host tissue *S* gets infected by different components of the pathogen population, at different rates when the fungicide is present. We assume that the different strains behave identically in the absence of control (i.e. we neglect modelling fitness costs).

Pathogen mutation is considered, so a small proportion of pathogen offspring has a different trait value to its parent. We use a Gaussian mutation kernel with mutation scale *σ*^2^ and assume mutation events occur with probability *p*. For a strain *j*, we describe the probability *P* (*k, j*) of its offspring taking trait value *k*:

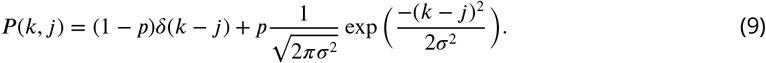

Here *δ*(*x*) is the Kronecker delta (which is 1 when *x* = 0 and 0 otherwise). All strains must have trait values within [0, 1], so any offspring that are predicted to take negative values are given trait value 0 and any that are predicted to take values greater than 1 are given trait value 1.

The per capita growth rate of offspring with trait value *k* is given by the integral over all possible parents with trait value *j*:

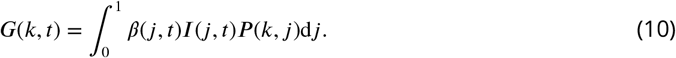

The total per capita growth rate of new infection at time *t* is given by the integral over all new strains:

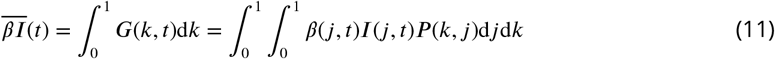

This means fastest growing strains have relatively higher contribution to new infections.

The *S* equation becomes:

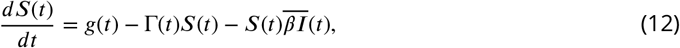

The *I* equation, for each strain with trait value *k*, becomes:

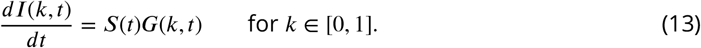

The total amount of initial inoculum is fixed at the same value each year, although evolutionary changes mean that its strain composition varies over years. This choice means the integral over all pathogen strains equals *I*_0_ at the start of each modelled season (*t* = *T*_1_, Table 2).

#### Incorporating host plant resistance (full model)

A similar modelling approach can be used to explore how the pathogen population adapts to resistant cultivars. We introduce a ‘host trait’ *l*. The main difference between the host trait and the fungicide trait in the model is that the host protection is assumed to act in the same way throughout the season (no time-variation), whereas fungicide control depends on the fungicide concentration which depends on time. The value of *k* is as before, whereas the value of *l* corresponds to the host effect at any time:

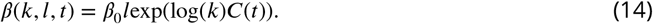

Higher host trait values give higher infection rates corresponding to increased pathogen fitness (similarly to the fungicide trait values).

Mutation is assumed to act in a similar way to before, with both *k* and *l* changing, again on a [0, 1] scale. The per capita growth rate of the pathogen strain with fungicide trait *k* and host trait *l* is given by the integral over all possible parent strains with fungicide trait *j* and host trait *m*:

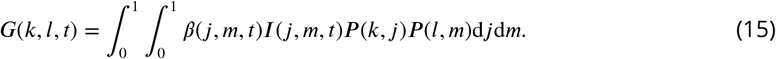

Host and fungicide mutations are assumed to act independently of each other.

The total per capita growth rate of infection is given by:

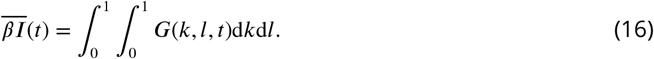

The model equations become:

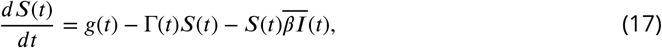

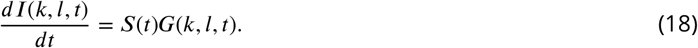

#### Notes on model implementation

When solving the model, we discretise the integral 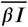 in Equations 12 and 17. We use at least 100 values for *k* (and *l* if using the full model) in each model run. The ‘discretisation number’ is specified in the parameter values in each figure caption, denoted by *n*_*k*_ for the number of *k* values (and *n*_*l*_ for the number of *l* values if applicable). We typically chose larger values for *n*_*k*_ when using the fungicide only model, since the full model was slower for large values of *n*_*k*_ and *n*_*l*_ (because the discretisation was then over a grid of *n*_*k*_ × *n*_*l*_ values rather than a one dimensional spectrum of *n*_*k*_ values).

The fungicide concentration equation (Equation 5) can be solved for each spray programme and written explicitly. For example, the concentration of a 3 spray programme is given by:

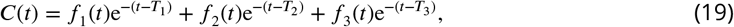

where

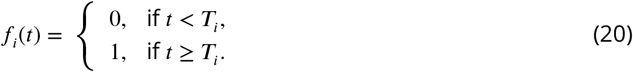

The form is similar for the 1 and 2 spray programmes, except the 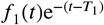 term is omitted for the 1 and 2 spray programmes and the 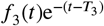 is omitted for the 1 spray programme. In our implementation of the model we use this form rather than solving the differential equation each time the model is run.

### Model fitting

#### Initial inoculum

We fitted the initial level of inoculum at the start of each season using Dataset A (see Table 3 and also Appendix 1). We assumed this initial level was constant over seasons, with differences in disease pressure from season to season captured entirely by the infection rate (*β*_0_), which varies over seasons in our model. The fitted value of *I*_0_ was used for all model runs unless stated otherwise. Dataset A was collected in Denmark from seven locations across three years. The trials reported in Dataset A were conducted by Aarhus University and the knowledge centre for agriculture, SEGES.

**Table 3.**
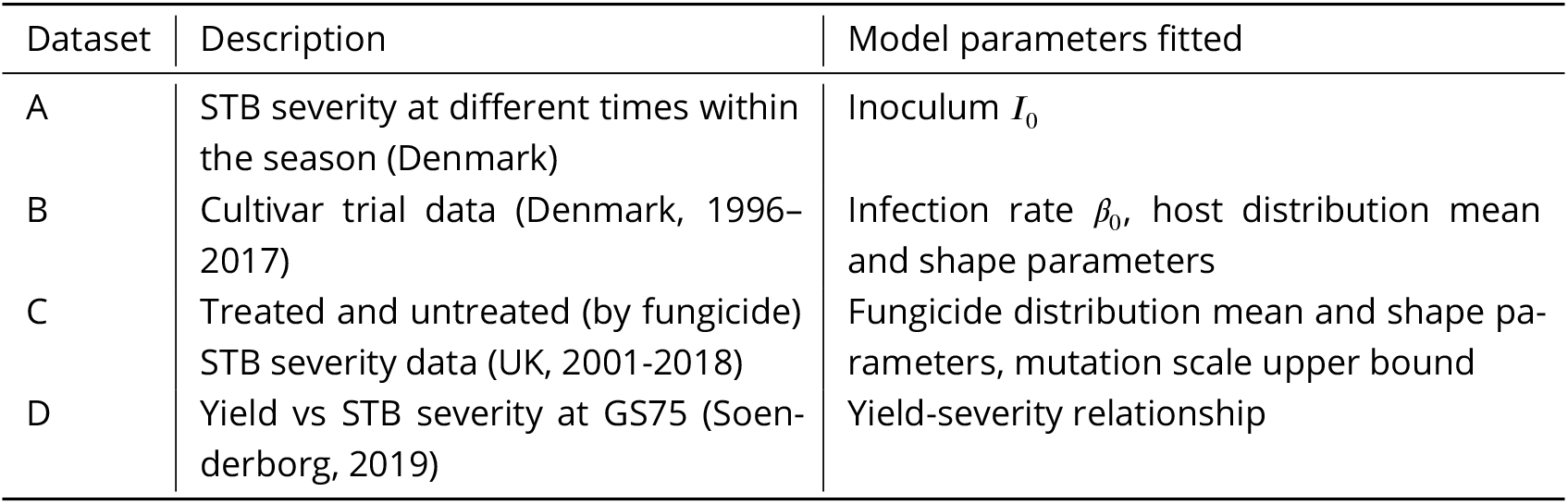
Datasets used in model fitting.

#### Infection rate

To find the distribution of infection rates *β*_0_, we used data from cultivar trials which were run from 1996 until 2017 (Dataset B). The trials from Dataset B were conducted by the knowledge centre for agriculture, SEGES and the Danish Institute of Technology. The data set includes data about disease severity at growth stage 75 on untreated plots, depending on location and cultivar variety. We filtered the raw trial data to consider plots that were untreated by fungicide and excluding those with cultivar mixtures. This is because we were interested in the level of disease in the absence of control measures. We also filtered to only include high pressure locations (higher than 6 on a 1-10 scale), because we were most interested in testing disease management strategies in the regions where disease is the most damaging.

We grouped the data by location, cultivar and year, before finding the mean severity score for each location/cultivar/year combination. For each year/location, we found the worst performing cultivar and assumed that this mean severity is representative of the average severity in the absence of control (i.e. that the pathogen has reached peak fitness for that cultivar). We chose ‘mariboss’ as the disease-resistant cultivar to analyse. It was one of only 4 cultivars which appeared for 10 or more years in the dataset, and it offered a relatively good level of control initially. We filtered the dataset to only include location/year combinations where mariboss was grown. All data for the worst performing cultivar (or cultivars if multiple cultivars had an equally poor average) in each year and location that mariboss was found were used as the values for severity in the absence of control.

We fitted a truncated exponential to these severity data, with a probability density function of the form:

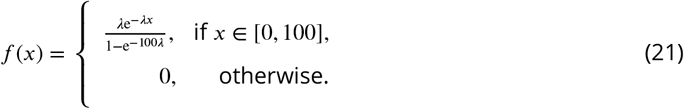

This allowed us to sample many values for final disease severities in the absence of control, which we used to generate infection rate (*β*_0_) values. This was done by numerically solving the ODEs using the value for *I*_0_ as found above. For each severity, we used a numerical optimiser (‘minimize’ from the open-source Python package ‘scipy.optimise’) to find the corresponding value of *β*_0_.

#### Mutation

The mutation kernel has two parameters, one for the proportion of offspring which have a different trait value and one for the scale of mutation, which affects how different these trait values are (Equation 9). We assume that the proportion and scales of mutation are the same for fungicide and host traits.

The value for the mutation proportion we use is based on estimates for the total number of spores generated in a growing season and the number of spores per season carrying adaptive mutations for fungicides and resistant cultivars (***McDonald et al., 2022***). See Appendix 2.

To identify an appropriate value for the mutation scale, we first find an upper bound for it. In general, the loss in control efficacy is caused by a combination of selection on strains that are already present in the population as well as on strains that arise due to mutation. This means we can find an upper bound from the value for mutation scale for which the loss in efficacy could be attributed to mutation alone with no initial standing variation (Appendix 2). We then arbitrarily choose 10% of this value to the be the value for the mutation scale used in the model, but we test this assumption in Appendix 3 and find that the model output is very similar for a wide range of choices of mutation scale.

#### Fungicide trait distribution

We fitted an initial distribution for the fungicide trait to reflect fact there is some initial variation in pathogen fitness relative to the fungicide. To find this distribution we used information derived from disease severity data (Dataset C) from UK disease trials from 2001-2018 (***van den Bosch et al., 2020***; ***Blake et al., 2018***). ***van den Bosch et al. (2020***) fitted curves to these data, of the form

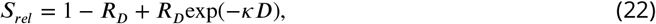

to 5 different dosages *D*, where *S*_*rel*_ is related to control by control = 100 − 100*S*_*rel*_, so that control takes values between 0% and 100%. They reported the mean value and standard error for the two parameters *R*_*D*_ and *κ* in each year. The parameter *R*_*D*_ relates to the maximum reduction in severity at theoretical infinite dose and *κ* is a curvature parameter.

We used a t-distribution (***Ruckstuhl, 2010***) with 3 degrees of freedom to drive a procedure based on the reported standard errors to sample 500 values per year for *R*_*D*_ and *κ* and then used these values to generate 500 control values each year. We filtered out any generated control values less than 0 and greater than 100 and resampled to replace them until we had 500 values in every year. The bias introduced by this filtering process was small, changing the mean value by 0.83% on average and 2.3% at most across the 18 years in the dataset. Using the uncertainty rather than just a single fitted value per year allowed a clearer assessment of how the model performed on the training and the test set given the variability in the data, and meant that years with lower uncertainty would have a larger influence on the optimal model parameters found in the model fitting process.

We use a gamma distribution to characterise the initial distribution of fungicide curvature parameter (*θ*, equivalent to −log(*k*)). The gamma distribution is a continuous, two parameter distribution (we use the shape and rate parameterisation) with support on (0, *∞*). This allows us to compare the mean of the fitted distribution for *θ* with those found in the literature in ***Hobbelen et al. (2013***); ***Elderfield et al. (2018***); ***Taylor and Cunniffe*** (***2022***). We found the optimal values for the two-parameter gamma distribution (i.e. the values that minimised the sum of squared distance between the data and the model output) using the open-source Python package Optuna as an optimiser.

The default untreated disease severity reported by ***van den Bosch et al. (2020***) was 37%, corresponding to the top 10% of disease severities measured in untreated plots, averaged across all years of trials. ***van den Bosch et al. (2020***) did not report a separate value for each year nor standard error terms, so we used this single value for the disease severity in the absence of control in each year. The cultivars used in the experiment were selected for their susceptibility to septoria (***van den Bosch et al., 2020***), so we assume that they offer negligable levels of control. We used an infection rate (*β*) value corresponding to the 37% value for every year.

We split our data into a training and a test set, using 2/3 of the data by year for the training set (the training set was from 2001-2012, and the test set from 2013-2018).

#### Host trait distribution

In each year and location (Dataset C), we found the level of control offered by the disease-resistant cultivar mariboss relative to the worst performing cultivar, using the following formula:

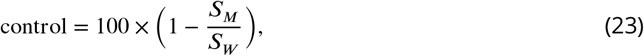

where *S*_*M*_ is the mean severity for mariboss and *S*_*W*_ is the mean severity for the worst cultivar in that particular year and location.

We use a beta distribution to describe the initial distribution of host trait values *l*. Beta distributions are continuous, two parameter distributions defined between the values of 0 and 1. They take a wide variety of shapes and commonly appear in population genetics contexts (***Balding and Nichols, 1995***). The host trait value *l* is defined on this range and so the beta distribution was an appropriate choice (in contrast to the fungicide trait parameter, for which a positive value was the only constraint since we fit a distribution of curvature values *θ* rather than trait values *k*).

Again we use Optuna to optimise the values of the two distribution parameters, resulting in the best fit according to a loss function *L* (Equation 24). Let *M*_*y*_ be the control prediction from the model with no fungicide applications in year *y*, and *C*_*y,l*_ be the cultivar control observed in the data in location *l* (since we have multiple locations in each year, up to *N*_*y*_ in year *y*). The control measurement *C*_*y,l*_ was determined from the relative severities of mariboss compared to the worst performing cultivar in each year/location combination that mariboss was grown in (Equation 23). Then let *n*_*y,l*_ be the minimum of the number of trials from the worst performing cultivar and of mariboss in the each year/location combination.

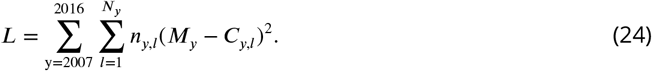

Again we separate the data into a training and test set with a 2/3 split by year (the training set was 2006-2012, and the test set was 2013-2016).

#### Yield

To analyse the relationship between disease severity and yield, we used trial data from Soenderborg (Dataset D), a high disease pressure region of southern Denmark. The data were collected in 2019. Although this dataset corresponds to trials from Soenderborg, we assume that the response of wheat yields to a particular level of disease is the same regardless of location. We used a generalised additive model (GAM) for the yield-severity relationship. We used the open-source Python package ‘pyGAM’ to fit a monotonically decreasing GAM with 5 splines. The choice of 5 splines was arbitrary, but chosen so that there were enough splines to fit the data well but not so many that the model was overfitting. See Supplementary Data 1 for a CSV file relating disease severity values to yield values.

## Results

### Basic behaviour of the model of fungicide resistance

Initially we consider the case where control is offered by the fungicide only, and a susceptible cultivar is used. During the season, the disease severity (total level of disease) increases. Different strains within the pathogen population grow at different rates depending on their fitness relative to the fungicide (Figure 1A). The increase of disease is approximately logistic when the fungicide is not present (Figure 1B). Strains with greater fitness relative to the fungicide are slowed to a lesser extent by each fungicide application.

**Figure 1.**
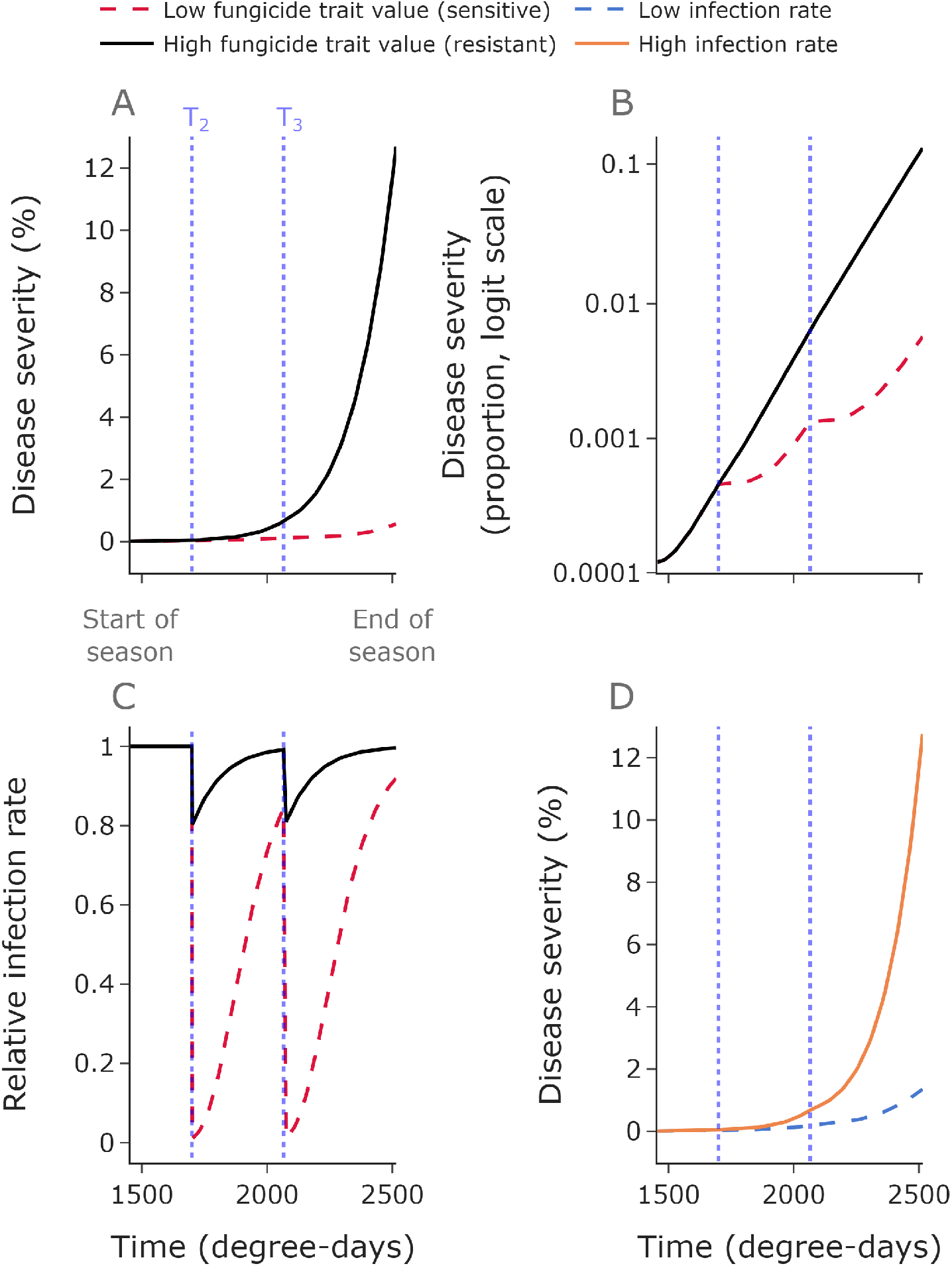
Disease severity depends on fungicide trait value and infection rate. The dotted blue lines show the two fungicide application times used here. The disease severity is much greater for a pathogen strain with a high fungicide trait value than for one with a low fungicide trait value **A,B**. Panel **B** shows the same disease progress curves on a logit scale (if *p ∆* (0, 1) is the disease severity, then its logit is logit(*p*) = log_10_(*p/*1 − *p*)), and shows how disease progression is approximately logistic (i.e. linear on a logit scale). Panel **C** shows how the relative infection rate varies depending on the fungicide trait values and as the fungicide concentration decays following applications. The baseline infection rate *β*_0_ affects the disease progress curve; high *β*_0_ values give greater disease severities than low *β*_0_ values. **Parameter values: A,B,C**: *β*_0_ = 0.008 degree-days^-1^, *I*_0_ = 0.01, low trait value = 0.01, low trait value = 0.8. **D**: low *β*_0_ = 0.006 degree-days^-1^, high *β*_0_ = 0.009 degree-days^-1^, *I*_0_ = 0.01, trait value = 0.4. The mutation proportion in all panels is 0. Here *n*_*k*_ = 100.

When present, the fungicide acts to reduce the infection rate by an amount which depends on the pathogen strain (Figure 1C) and on the concentration of chemical, which decays exponentially after each fungicide application. Disease pressure varies by year; this can be represented in the model by changing the value of the infection rate *β*_0_ in each simulated year (Figure 1D).

### Model fitting

The truncated exponential fits the disease severity data and was used to sample many values for final disease severities (Figure 2A, Table 4). The infection rate (*β*_0_) values generated from the observed data are plotted in Figure 2B, along with the probability density function for *β*_0_ generated from the truncated exponential distribution for disease severities. The result of the fitted GAM used for the yield-severity relationship was a smoothly varying curve through the data (Figure 2C). The optimal model fit, including mutation parameters and initial trait distribution leads to a control curve passing through the data, including extrapolation to ‘test’ data not used as ‘training’ data for fitting the model (i.e. years past 2012, Figure 2D).

**Figure 2.**
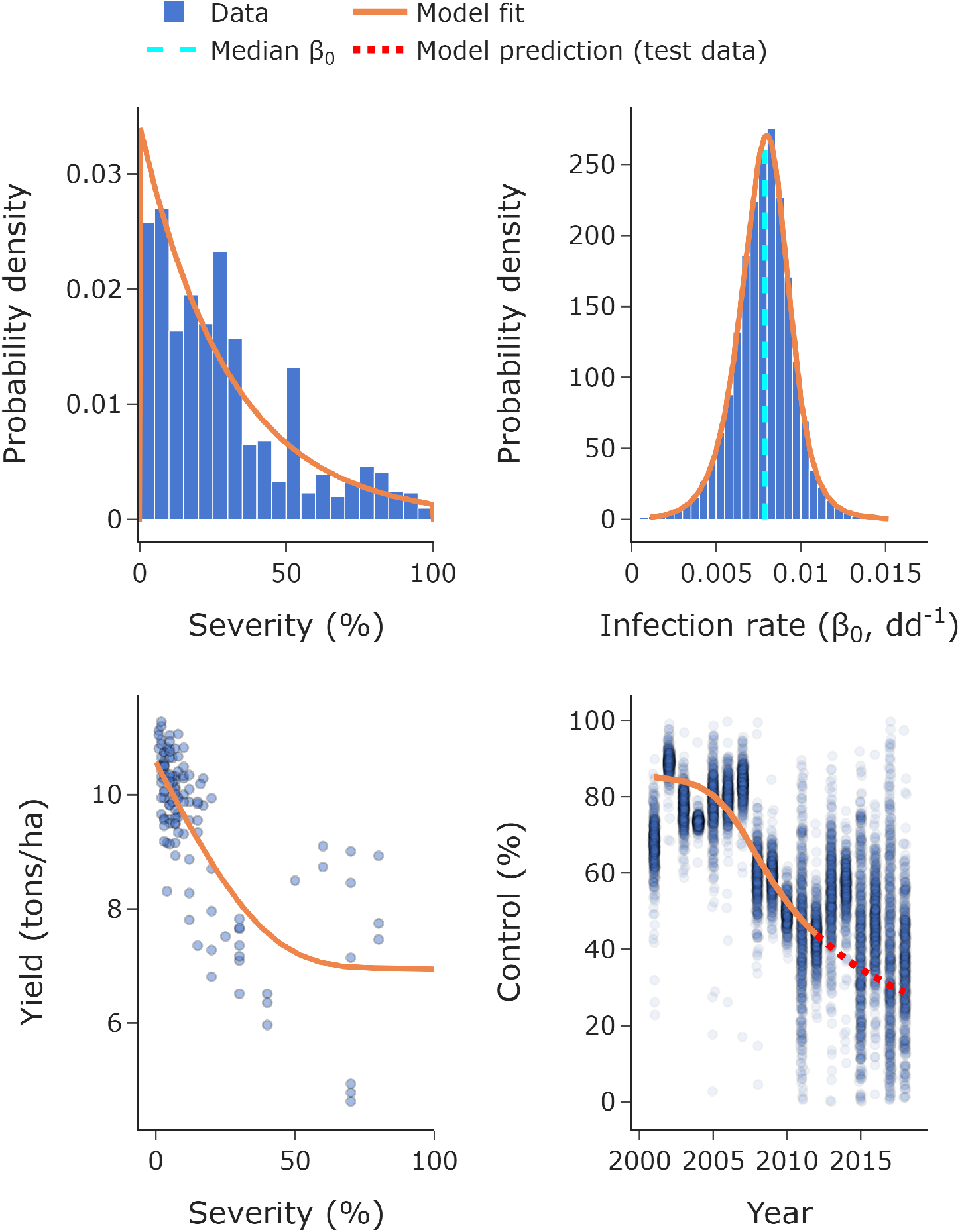
Model fitting. We fitted a truncated exponential to the severity data, allowing us to sample as many random values for disease severity as we need for each simulation (**A**). From these severities and the fitted value for inoculum *I*_0_ we can find a probability density function for the infection rate *β*_0_, shown here alongside a histogram of values inferred from the observed severity data (**B**). Here ‘dd^-1^’ is degree-days^-1^. We used a generalised additive model to describe the relationship between yield and severity (**C**). We used fungicide control data from 2001 to 2012 (solid line) to find the optimal initial trait distribution parameters and used data from 2013 to 2018 (dotted line) as a test set **D**. Here *n*_*k*_ = 400.

**Table 4.** Parameter values used in the model (to 3 significant figures). The parameter values are fitted to data. The sampled infection rates are generated from a truncated exponential distribution with maximum 100 and parameter *λ* (see Equation 21).

| Parameter | Symbol | Range/Value | Dataset/citation | Figure |
| --- | --- | --- | --- | --- |
| Initial inoculum | $I_0$ | 0.00986 | A | Appendix 1 Figure 1 |
| STB truncated exponential parameter | $\lambda$ | 0.0329 | B | Figure 2A |
| Infection rate | $\beta_0$ | Sampled | B | Figure 2B |
| Median infection rate | $\beta_{0,M}$ | 0.00787 | B | Figure 2B |
| Mutation proportion | $p_M$ | $1.23 \times 10^{-5}$ | *McDonald et al. (2022)* | - |
| Mutation scale, upper bound | $\sigma^2$ | 0.0198 | C | Appendix 2 Figure 1 |
| Fungicide distribution mean | $\mu_F$ | 9.44 | C | Figure 2D |
| Fungicide distribution scale parameter | $s_F$ | 1.19 | C | Figure 2D |
| Host distribution mean | $\mu_H$ | 0.809 | B | Figure 5A |
| Host distribution mean shape parameter | $b_H$ | 6.59 | B | Figure 5A |

### Model outputs

#### Comparing numbers of fungicide applications

We explore the effect of the number of fungicide applications per year on the pathogen population, disease severity and yield. The results shown in Figures 3 and 4 do not include disease-resistant cultivar control and consider the effect of fungicide applications only. Figure 3 show the average result across an ensemble of runs, whereas Figure 4 shows the variability in outcomes from the ensemble.

**Figure 3.**
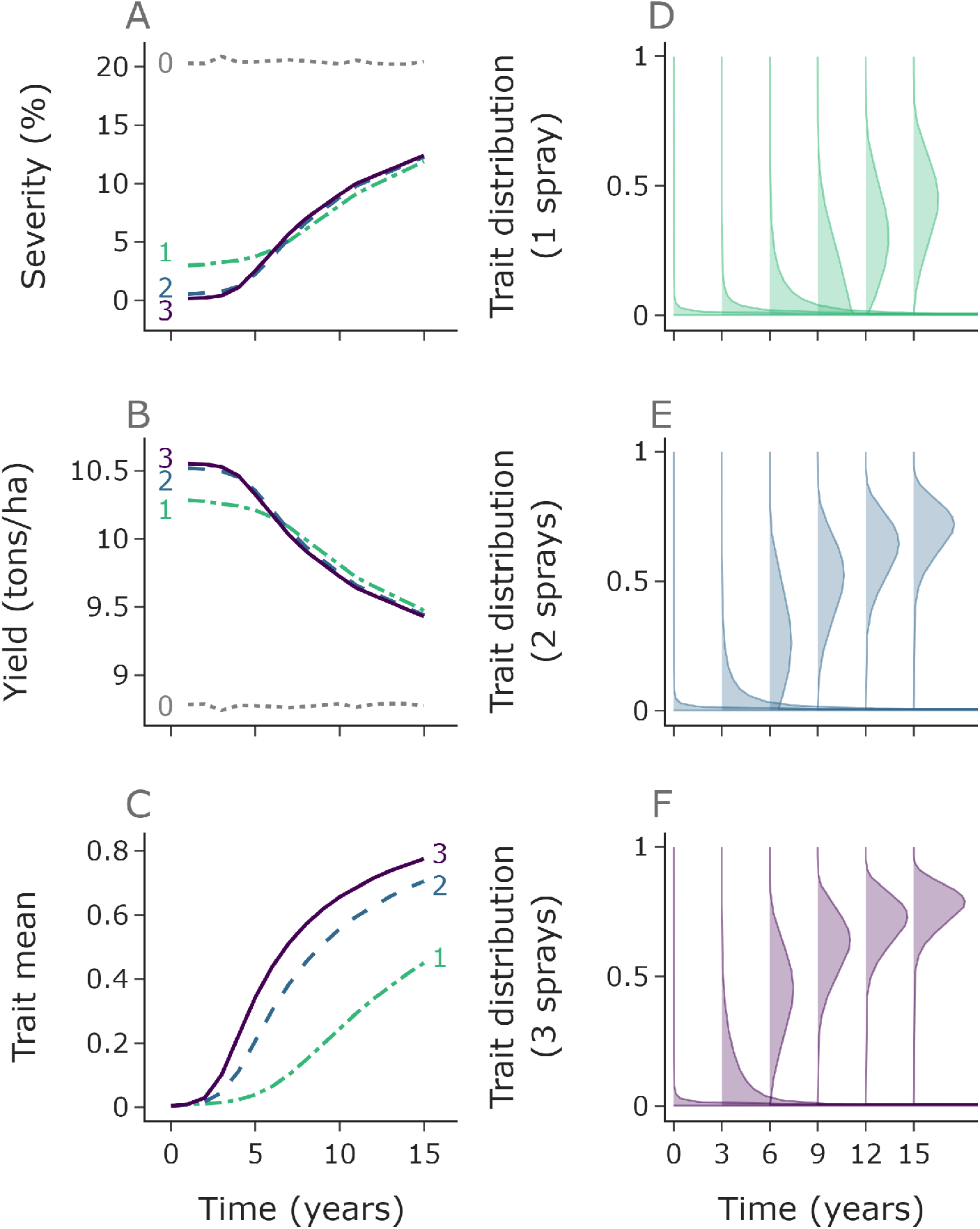
More fungicide applications per year leads to faster selection for resistant strains and to lower yield in the long term. Median values for disease severity and yield from each year of the 10,000 simulations (**A, B**). Initially three fungicide applications per year gives the lowest severity and highest yield. Panels **C**-**F** relate to the ‘mean distribution’ across the 10,000 ensembles, and the mean value of this mean distribution in **C**. Due to more rapid selection for resistant strains with more fungicide applications (**C**-**F**), the one fungicide application per year strategy outperforms the two and three applications per year strategies in terms of both severity and yield by year 7. Here *n*_*k*_ = 500.

**Figure 4.**
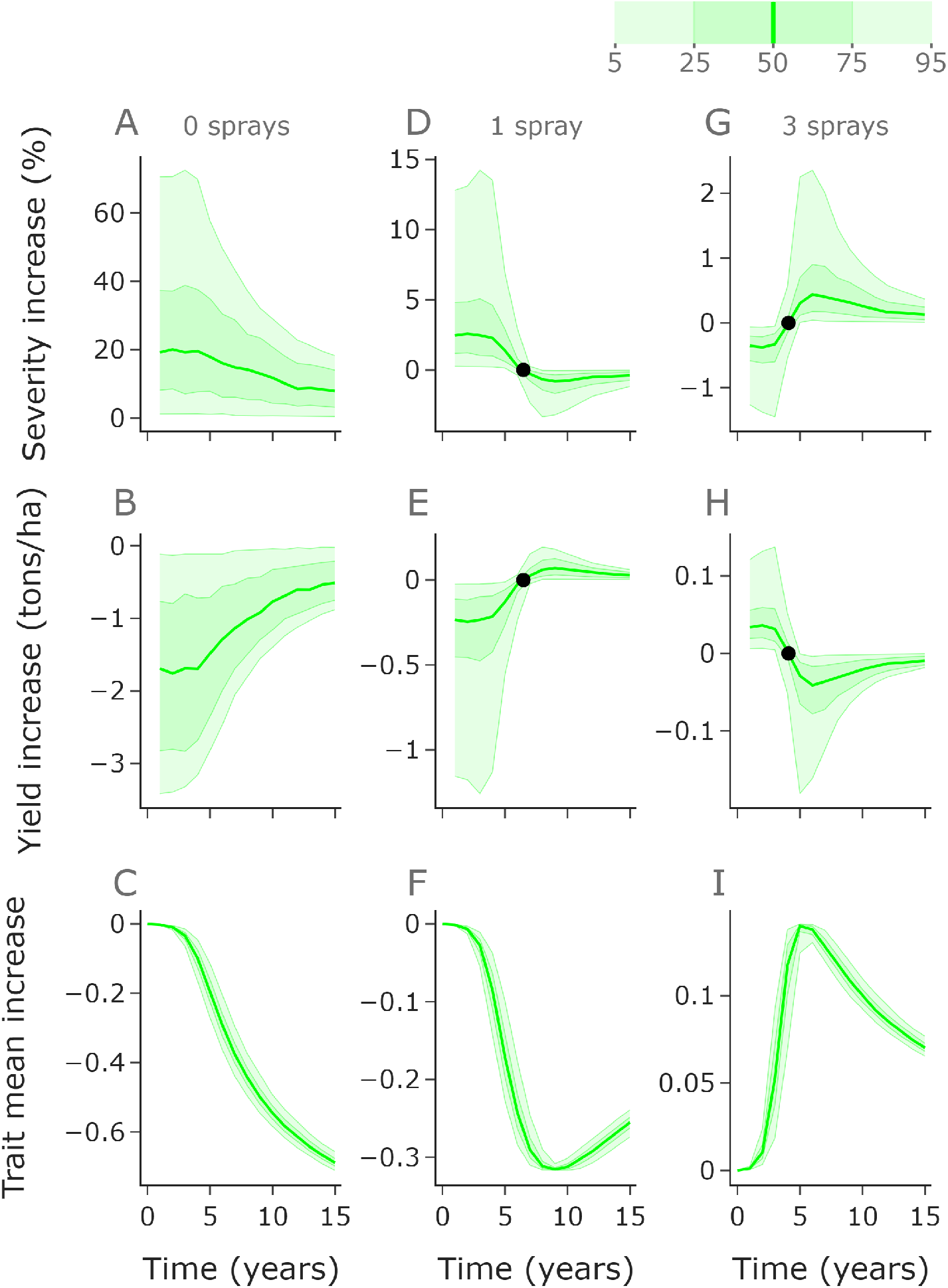
The optimal disease management strategy depends on the relevant timeframe. In the left, middle and right columns we compare the performance of zero, one and three fungicide applications annually with the performance of two applications annually. Zero applications (no control) leads to higher severity (**A**) and lower yield (**B**) than two applications, although the difference reduces over time due to selection for resistant strains (**C**). Initially one application leads to increased severity (**D**) and decreased yield (**E**), but by year 7 one application is better than two due to reduced selection for resistant strains (**F**). Conversely, three fungicide applications initially leads to lower severity (**G**) and higher yield (**H**), but by year 5 this benefit is lost due to increased resistance (**I**). Here *n*_*k*_ = 400.

Initially, a greater number of applications offers greater control than fewer (Figure 3A). The figure shows the median value for disease severity and yield in each year over an ensemble of 10,000 runs, each one using randomly sampled values (from our fitted distribution) for the infection rate *β*_0_. It shows the mean density of the distribution at each discretised value for *k*, resulting in an ‘average’ distribution across the ensemble (Figure 3C,D,E,F). The strategies with more fungicide applications lead to faster selection for higher trait values, leading to a gradual shift in distribution (Figure 3D,E,F). This faster shift in distribution leads to the order in terms of disease severity and yield reversing such that 1 fungicide application per year outperforms 2 and 3 applications per year (Figure 3A,B). The higher the number of applications, the more rapidly the distributions shift, which can be shown most clearly by looking at the mean of the distributions (Figure 3C).

Initially the control offered by just one application is strong. This means that the improvement in severity and yield offered by further applications is marginal, with two and three applications performing comparably initially. Their performance remains similar in subsequent years due to a trade off – more applications means greater amount of time with fungicide control, but also faster selection for resistance. This trade off leads to two and three strategies behaving similarly, whereas by year 7 the reduced selection from the one fungicide application strategy outweighs the reduced control caused by fewer applications, and after year 7 the single fungicide application programme performs optimally for yield.

#### Ranges of outcomes under different numbers of fungicide applications

To further explore the relative performance of differing numbers of fungicide applications, we compare the full distribution of outcomes under zero, one and three applications per year to two applications (Figure 4). We use 1,000 simulations with infection rates sampled randomly from our fitted distribution from Figure 2B. The same set of sampled infection rates were used for each application strategy.

Zero applications per year (no treatment) gives higher severity and lower yield (Figure 4A,B) than two applications per year, with particularly extreme differences in the early years whilst the fungicide still offers good control in the two application programme. Within 15 years the fungicide loses efficacy due to the shift in pathogen distribution (Figure 4C), meaning that the control offered is reduced.

One application per year initially gives higher severity and lower yield (Figure 4D,E) than two application per year, but by year 7 the median yield is higher and severity is lower due to two applications leading to higher levels of resistance (Figure 4F), as indicated by the summarised results in Figure 3.

Three applications per year initially reduces severity and increases yield relative to two (Figure 4G,H). However, by year 5 the increased resistance in the pathogen population (Figure 4I) leads to two applications outperforming three. The differences in performance are less exaggerated for the three versus two comparison than for the one versus two or zero versus two comparisons. This is because the difference in control (at least initially) between two and three sprays is smaller than between one and two, which is smaller than between zero and two.

#### Introducing disease-resistant cultivars

Figure 5 shows the results of introducing disease-resistant cultivar control, comparing the results of a two-spray programme with and without host-plant protection from the disease-resistant cultivar mariboss. The model fit for the initial trait parameters that optimally matched the decline in control offered by mariboss in the absence of fungicide treatment is shown in Figure 5A. The disease-resistant cultivar helps reduce disease severity and increase yield (Figure 5B,C). The use of the resistant host variety reduces the disease pressure, which protects the fungicide, as shown by the difference in fungicide trait mean between the strategies (Figure 5D). This protection offered by the host to the fungicide means that although the difference in yield and severity in year 1 is small, by year 7 the improvement in yield is much larger, particularly in high disease pressure years. This is because the control offered by the fungicide alone is excellent initially, so both strategies offer excellent control and the difference in yield is minimal. However, in later years the benefit of choosing a resistant cultivar becomes significant – greater than the initial benefit of using 2 sprays rather than 1 or 3 rather than 2 in Figure 4E,H.

**Figure 5.**
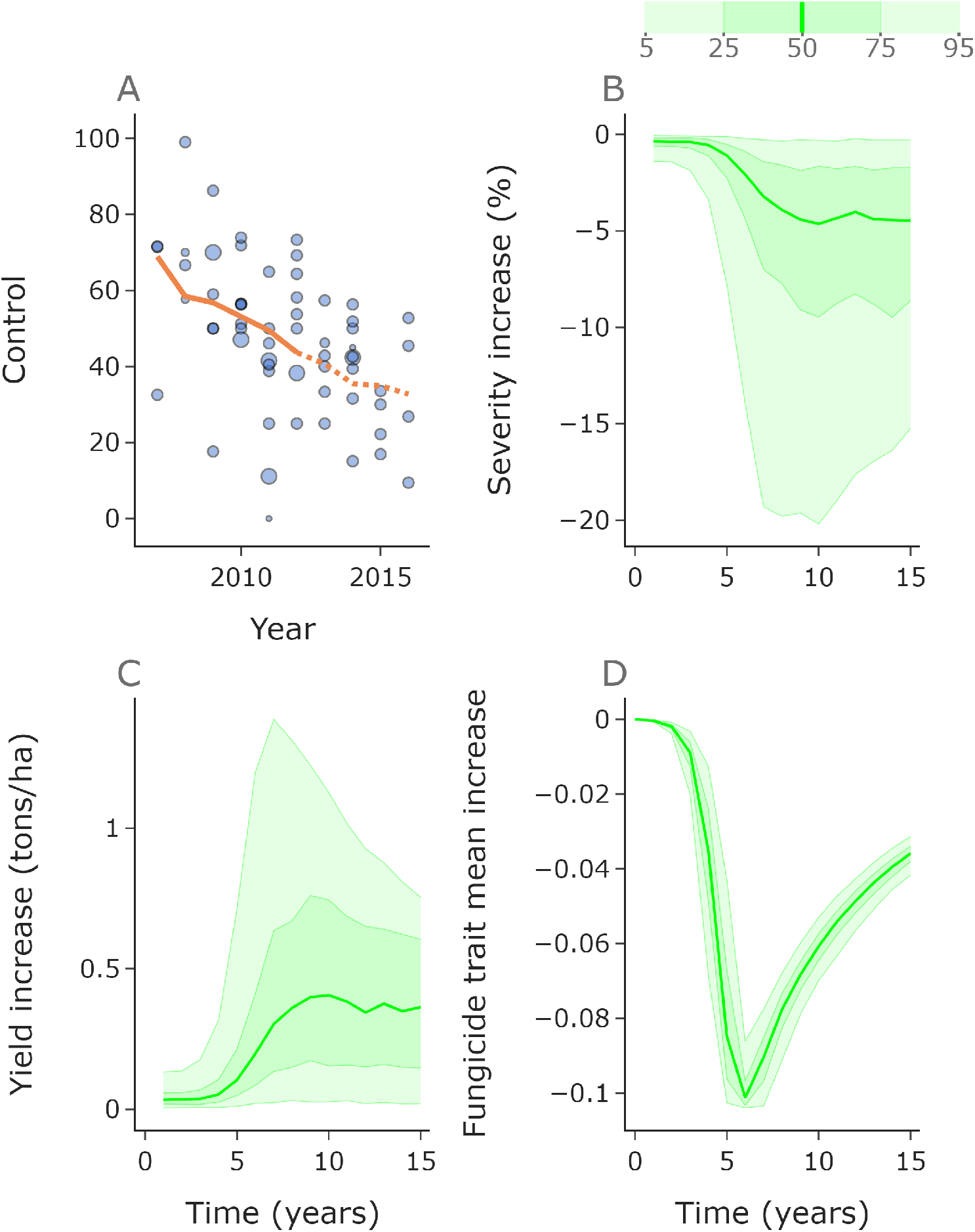
Disease-resistant cultivars reduce severity and protect the fungicide. The optimal host trait initial distribution parameters were fitted using data from 2007 to 2012 (solid line, **A**). We used data from 2013 to 2016 as a test set (dotted line). When comparing a two application fungicide programme with and without the disease-resistant cultivar, the disease-resistant cultivar reduces the severity (**B**) and increases yield (**C**); in each case the response shown is the difference between two spray fungicide programmes with and without the resistant cultivar. Further, the fungicide develops resistance more slowly due to reduced disease pressure when the resistant cultivar is used (**D**). This means the benefit of using the resistant cultivar becomes greater as time goes on, and is particularly noticeable in high disease pressure years with yield improvements of up to 1.39 tons/ha in the 95th percentile in year 7. Here *n*_*l*_, *n*_*k*_ = 100.

#### Varying host control and regular replacement of disease-resistant cultivars

To further explore the extent to which a disease-resistant cultivar protects the fungicide (and vice versa), we ran a parameter scan testing the performance of different hypothetical cultivars. We allowed the initial mean of the host trait distribution to vary whilst keeping the shape parameter of the distribution fixed at the value from the model fitting. Some examples of the resulting distributions are shown in Appendix 4. For simplicity, we used the median value *β*_0_ = *β*_0,*M*_ (Table 4) in all examples, i.e. the underlying simulations did not account for environmental stochasticity.

For very short timescales (5 years), more fungicide applications gives improved yield (Figure 6A). However over longer timescales more applications leads to reduced yield for all but the strongest of our theoretical host cultivars (Figure 6B). This is because selection for fungicide resistance is stronger with more applications, although for particularly strong host protection higher numbers of fungicide applications are preferable due to increased mutual protection between host and fungicide. It should be noted that these particularly strong hosts have vastly stronger protection than the fitted cultivar mariboss (dotted vertical line).

**Figure 6.**
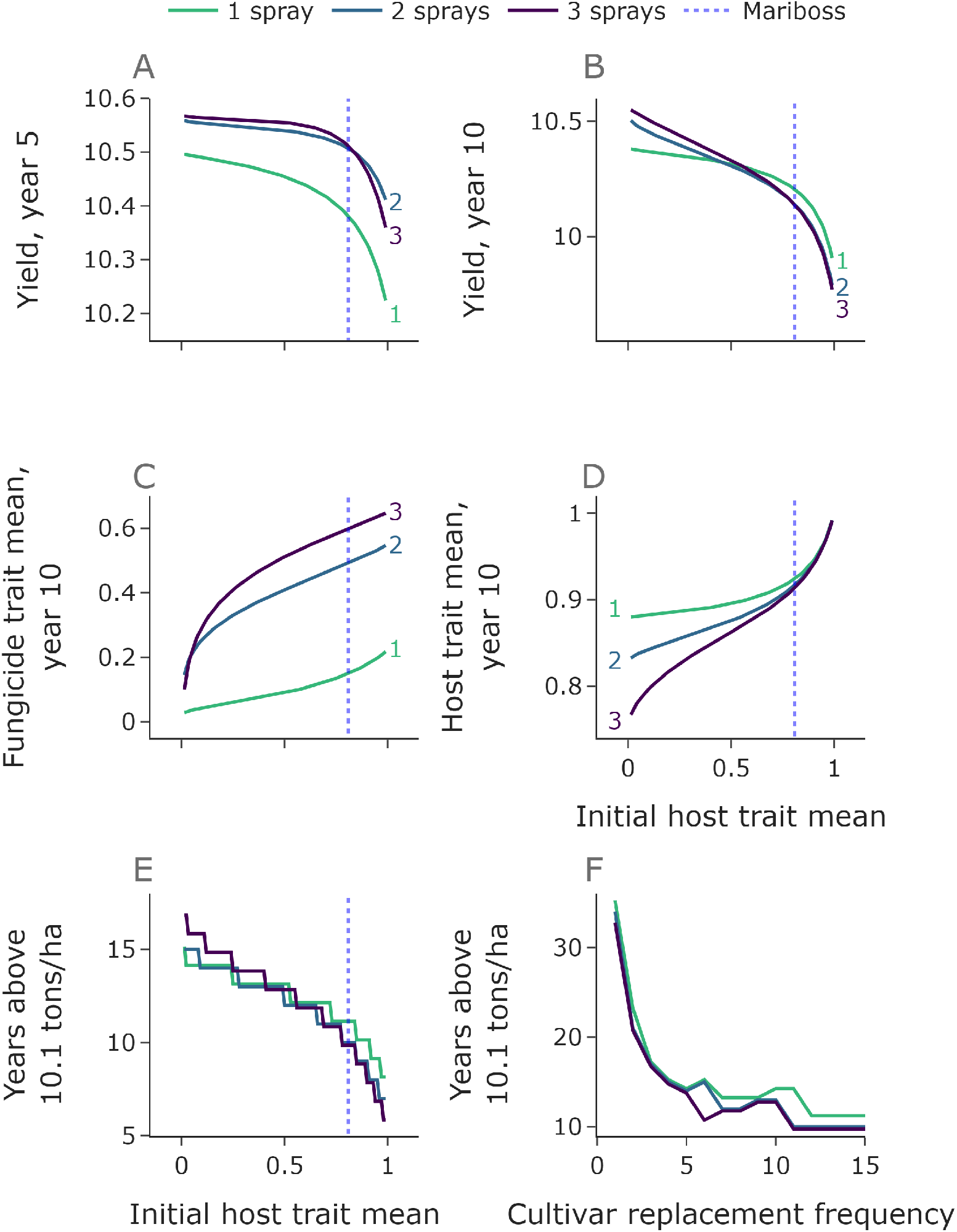
More effective cultivars protect fungicides more strongly and more frequent replacement leads to effective control for longer. In panels **A**-**E** we vary the initial host trait mean whilst keeping the shape parameter fixed. The vertical dotted lines show the fitted trait mean value for mariboss. Lower host trait means than this value represent hypothetical cultivars which are even more disease-resistant. Over a short timescale (5 years) more applications is preferable than only one (**A**). However, for longer timescales and initial trait values close to that of mariboss (dotted line) one application outperforms two and three. For a given number of fungicide applications per year, the stronger the host protection (lower trait values), the slower the fungicide resistance develops (**C**). For a given initial host strength, higher numbers of fungicide applications protects the host more effectively (**D**). One application keeps the yield above a threshold of 10.1 tons/ha for the longest for all initial host trait values greater than 0.56 (**E**). Each year value is an integer but we offset each line vertically by a small amount to show the ordering. If cultivars are replaced regularly then the yield remains above the threshold for much longer (**F**). In this panel the cultivars are assumed to have identical performance to mariboss. Here *n*_*l*_, *n*_*k*_ = 100.

Strategies with more fungicide applications lead to faster selection for resistant strains and correspondingly a higher fungicide trait mean in year 10 (Figure 6C). Stronger host protection (lower initial host trait mean) leads to lower values of the fungicide trait mean due to the protection offered by the cultivar to the fungicide via reducing disease pressure. Similarly, the host trait mean after 10 years of the integrated management strategy is reduced when more fungicide applications are used (Figure 6D), since more intensive fungicide application programmes offer stronger protection to the host. This difference is less exaggerated for cultivars with efficacy comparable to mariboss than for hypothetical ones with extremely low initial trait means.

As an overall summary of the durability of the combined programme, we compared the number of years for which yield remains above a threshold of 10.1 tons/ha. This threshold value was chosen as a value slightly higher than the yield achieved with a resistant cultivar but no fungicide input (9.98 tons/ha). One application outperforms two and three for all host trait values above 0.56 (Figure 6E), although note that this result depends on the choice of threshold (for very high yield thresholds one fungicide application is not sufficient to achieve a high enough yield even in year one).

We ran a second scan where we assume that the cultivar is replaced at regular intervals, from every year to every *N* years (Figure 6F). In practice this would correspond to wheat breeders releasing new varieties to market. Each time the current cultivar is replaced with another cultivar with efficacy matching that of mariboss, but to which the pathogen population has evolved no resistance. This prolongs the effectiveness of the host protection and increases the protective effect offered by the host to the fungicide. For very regular replacement (replacement frequency 1 or 2), the number of years above the threshold is much higher than without replacement. The jagged nature of the lines is caused by the hard 10.1 threshold – for example, there is a sharp jump between 10 years and 11 years for 3 sprays, because without any replacement the strategy would fail in 10 years but if replacement happens in the final (10th) year then the sudden improvement in cultivar control leads to several more years of acceptable yields. With a replacement frequency of 5 years we get replacement in the 5th and 10th years before failure, but with a replacement frequency of 6 years the strategy fails in the 11th year just before the second replacement, so the outcome is significantly worse.

## Discussion

Integrated pathogen management strategies will be required for sustainable control of STB (***McDonald and Mundt, 2016***). Many such strategies involve combining chemical control with resistant cultivars. Pathogen evolution poses a severe threat to the effectiveness of fungicide and cultivar control of crop diseases, meaning that understanding how to optimally combine these disease management strategies will be crucial to maintaining strong crop yields. Although many fungicides currently in use are challenged by quantitative pathogen resistance (for example the azoles), the model presented here is the first model of quantitative fungicide resistance to be fitted to field data (Figure 2). Further, the model incorporates both cultivar and fungicide control.

Our analysis shows that more fungicide applications leads to higher levels of resistance in the pathogen population (Figure 3). However, for a given level of resistance in the population, more fungicide applications gives more control. This trade-off affects which application strategy is optimal. Over short time frames, high numbers of fungicide applications lead to improved disease control. However, over longer timescales reduced numbers of applications are preferable since they reduce the selection for resistant strains and eventually this leads to higher yields. Another consideration is how risk-averse the grower is – in particularly high disease pressure years the improved control offered by higher numbers of application can be notable (Figure 4). One approach could be to use more applications in the highest disease pressure years but in general use the minimum number of applications possible to achieve a desired level of disease control, perhaps with the aid of a decision support system (***Lázaro et al., 2021***).

The model shows that pathogen control offered by deployment of disease-resistant cultivars is an effective way to reduce disease severity and increase yield (Figure 5). There is an added benefit in using resistant cultivars that it delays fungicide resistance development (Figure 5D). This is caused by the cultivar reducing the pathogen infection rate. By reducing the infection rates of all strains, the difference between per capita infection rates is reduced, thus reducing selection according to the so-called ‘governing principles’ (***van den Bosch et al., 2014***). Similarly, fungicide applications can delay loss of cultivar control, in agreement with ***Carolan et al. (2017***). The more effective the cultivar, the more exaggerated the mutual protection effect (Figure 6). Again, over a shorter timescale higher yields are obtained with more fungicide applications per year, but over a longer timescale the outcome improves with fewer fungicide applications per year (Figure 6C,D). Frequent replacement of disease-resistant cultivars could be an effective way to maintain disease control (Figure 6G).

There are various modelling assumptions that were used for simplicity or due to a lack of available data. We neglect modelling fitness costs (***Hawkins and Fraaije, 2018***), since we did not have data to inform how different strains might carry fitness penalties. STB has a latent period where infection is asymptomatic (***Fones and Gurr, 2015***), which for simplicity is not modelled here. We also assume that the infection rate is fixed within each season – a more complex model could include temporal variation in infection rate within a single growing season, whereas we only change the infection rate value between different growing seasons. However, particularly when compared to the vast majority of fungicide resistance models (***Hobbelen et al., 2011a, 2013***; ***van den Berg et al., 2013***; ***Elderfield et al., 2018***; ***Taylor and Cunniffe, 2022***) in which environmental stochasticity is not considered at all, we contend this is a sensible treatment. We also follow many past modelling studies in using a fixed value for inoculum (***Hobbelen et al., 2011a, 2013***; ***Elderfield et al., 2018***; ***Taylor and Cunniffe, 2022***) but inoculum will vary year on year, and further the type of inoculum (ascospores vs pycnidiospores) can have complex effects on the latent period and the dynamics over multiple seasons (***Suffert and Thompson, 2018***). Finally spatial effects are ignored, despite promising early studies showing the potential of fungicide application patterns based on spatial risk (***Liu et al., 2017***). This theme of spatial deployment is particularly well established in the parallel host plant resistance literature (***Rimbaud et al., 2021***; ***Fabre et al., 2015***; ***Watkinson-Powell et al., 2020***). We have also implicitly assumed the pathogen strain composition in the inoculum is set by the grower’s own actions in previous seasons, which is equivalent to assuming a population of growers who all act identically and who all suffer the consequences of their actions. In practice, of course, there is more complexity and growers can be incentivised by economics to both prioritise or deprioritise resistance management (***Day et al., 2021***; ***Vyska et al., 2016***). Although ignoring these effects is a very common simplification in modelling studies, we note game theory (***Murray-Watson et al., 2022***) is an attractive framework to go beyond this.

The model fitting process was somewhat limited by the nature of the data available. Ideally we would have data on both the response of isolates to fungicide and cultivar control as well as the frequencies of these isolates in the pathogen population over many years. Further, to incorporate fitness costs into the model, we would need data on the relative infectivity of these isolates in the absence of cultivar control. For azole resistance, this may be difficult to characterise due to the complexity of the CYP51 fitness landscape (***Hawkins and Fraaije, 2021***). Although EC50 values have been recorded for different pathogen isolates (***ahd, 2020***), we did not have sufficiently many years of these data available from the same experimental setup to support the model fitting process. Further, the mapping from experimentally measured EC50 values of pathogen strains to the effect of the fungicide on those strains’ infection rates is not yet characterised. Although we were able to fit the model based on the decline in disease control, we had to infer the initial pathogen distribution and use the final few years of data for model validation rather than having data to describe the initial pathogen distribution and showing that this led to a decline in control comparable to that observed in the field. If we had access to data on the full distribution in successive years, the model fitting would become more powerful.

The model could be extended to address other disease management questions. Many of these future lines are obvious extensions now we have a fitted model of quantitative resistance in hand, given the long history and diverse applications of past models of mongenic resistance (***Hobbelen et al., 2011a, 2013***; ***Elderfield et al., 2018***; ***Taylor and Cunniffe, 2022***). We have considered full-dose applications of fungicide here; future work could include an analysis of dose choice and optimise for number of applications and dose in each application. A basic economic analysis could be included by accounting for the cost of each fungicide application (***Alves et al., 2021***; ***van den Bosch et al., 2020***). By extending the model to include two fungicides we could address questions on fungicide mixtures and whether mixtures outperform alternations in the case of quantitative resistance. We have considered strategies that are fixed in time, but it would be interesting to explore strategies that respond to the level of disease pressure and the level of resistance through time, e.g. something analogous to the so-called ‘Merry Dance’ (***Oliver, 2016***). There are many other crop diseases which are controlled by azole fungicides, for example powdery mildew, brown rust in wheat crops, citrus fruit mould and soybean rust (***Price et al., 2015***; ***Alves et al., 2021***). There are also pathogens (including *Z. tritici*) controlled by other fungicides to which resistance is quantitative, for example demethylation inhibitors (***Blake et al., 2018***). With appropriate data describing how yield of untreated plots compares to treated plots over many years, the model could be adapted to explore optimal management of these pathosystems, as well as accounting for combinations of chemical treatments. Many of these are questions that have been addressed, perhaps only partially, in the case of monogenic resistance (***Hobbelen et al., 2011a, 2013***; ***Elderfield et al., 2018***; ***van den Berg et al., 2013***), but for which the modelling framework here offers the possibility of transferring to a more complex, but more realistic, setting.

## Availability of code online

An implementation of the model in the freely-available programming language Python (Python Software Foundation, available at http://www.python.org) is online at https://github.com/nt409/ quantitative-resistance.

## Acknowledgements

We acknowledge SEGES and Tystoftefonden for their trial data used in fitting the model, and Rose Kristoffersen for useful discussions, particularly on interpretation of the data in the initial phase of model development. We thank Lise Jørgensen and Frank van den Bosch for useful discussions. NPT acknowledges the Biotechnology and Biological Sciences Research Council of the United Kingdom (BBSRC) for support via a University of Cambridge DTP PhD studentship (Project Reference 2119688). For the purpose of open access, the author has applied a Creative Commons Attribution (CC BY) licence to any Author Accepted Manuscript version arising from this submission.

## Appendix 1 Fitting initial inoculum

We did not have data to describe the level of inoculum (*I*_0_) at the beginning of the modelled season (*t*_*start*_ = *T*_1_), so we used disease severity data (Dataset A) for septoria severity from three time points (*t*_*A*_, *t*_*B*_ and *t*_*end*_, Table 2) within a single growing season to fit a value. We filtered the data so that we only considered plots untreated with fungicide, and trials for which the disease severity was greater than 0.

We used the freely-available Python package Optuna to sample potential values for the infection rate *β*_0_ and initial inoculum *I*_0_. For each pair of values, we found the squared distance between the logit disease severities from the model and from the data at each time point. The logit is defined as follows:

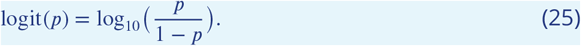

The logit scale was chosen since the disease progression is approximately linear on a logit scale and this choice means the severities from earlier in the year have a comparable impact on the model fit to those later in the year. The optimal parameters were the pair of values which minimised the sum of the squared distances (Appendix 1 Figure 1).

**Appendix 1 Figure 1.**
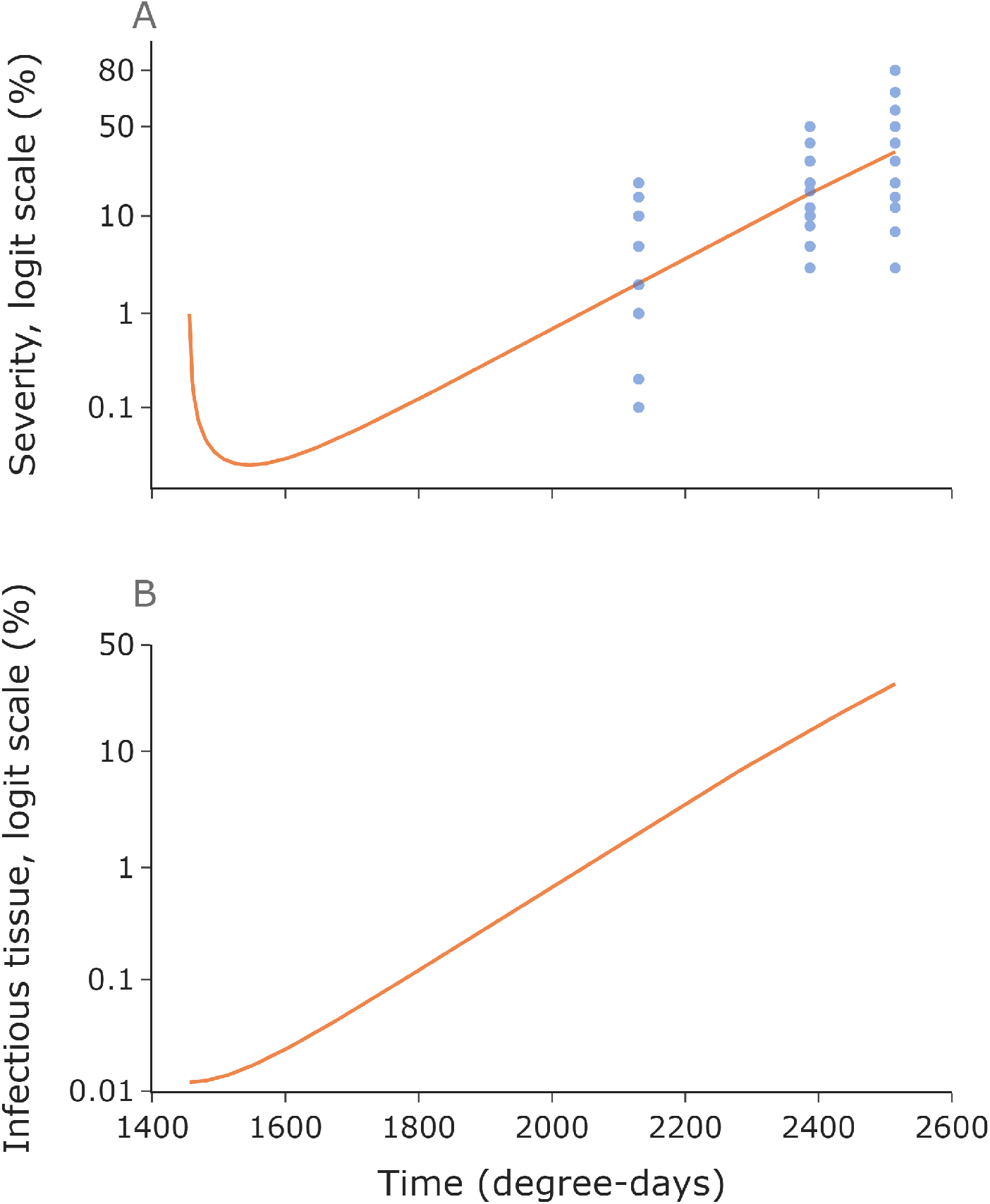
Fitting *I*_0_. We find the optimal values for initial inoculum *I*_0_ and infection rate *β*_0_ to fit the observed severity data. Initially the model severity decreases since the host grows faster than the infection (***Ferrandino, 2008***; ***Bailey et al., 2009***) (**A**). However, the amount of infectious tissue monotonically increases throughout the season (**B**). Note that infectious tissue is given as a percentage of the maximum amount of tissue after host growth is complete.

Instead of using the value of *β*_0_ found by this process, we had a much larger dataset with severities at the end of the modelled season (growth stage 75) which allowed us to find a full distribution of infection rates to sample from which facilitated an exploration of the effects of environmental stochasticity, as described in the main text.

## Appendix 2 Fitting mutation parameters

***McDonald et al. (2022***) estimate that wheat fields typically produce 2.3 to 10.5 trillion pycnidiospores per hectare, of which between 28 and 130 million spores carry adaptive mutations to counteract fungicides and/or resistant cultivars. This allows us to produce an estimate for the proportion of offspring which carry a mutation by dividing the average number of spores carrying mutations by the average total number of spores:

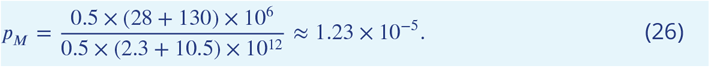

Finding a value for the mutation scale parameter was more complicated. We chose to consider the scenario with no standing variation, meaning that the initial pathogen population consisted of a single pathogen strain and mutation was the sole driver of loss of control. This gives us an upper bound for the mutation scale. In general we expect control breakdown to be caused by a combination of selection for existing resistant strains and well as on strains that arise due to mutation.

We used the same dataset for fungicide control as used in the fungicide fitting process described in the materials and methods section (Dataset C). We only considered the first and last years since we were seeking an upper bound for the mutation scale, meaning that the overall loss of control over the full timescale was more important than a close match to the shape of the control breakdown curve. The shape of the breakdown curve depends on the initial trait distribution, but we were most interested in showing that the model would lead to a sufficiently rapid loss of control for the appropriate choice of mutation scale. We used the fitted value from the fungicide control data (***van den Bosch et al., 2020***) directly rather than using the standard error values to generate distributions of values since we were only interested in the magnitude of the breakdown from the first to the final year.

We used the open-source Python package Optuna to find the optimal initial trait value and mutation scale (Appendix 2 Figure 1). The optimal parameter values were those that led to the minimal sum of squared residuals between the control values from the model and the data in 2001 and 2018 (first and last years).

For all other model runs we assumed that some of the breakdown of control was caused by initial standing variation in the pathogen population, and some was caused by mutation of the pathogen population. The mutation scale used in the model was 10% of the upper bound found in this section, arbitrarily chosen due to the lack of data to determine a value. The 10% assumption/choice is tested in Appendix 3. We also assumed that the host mutation scale is the same as the fungicide mutation scale. The mutation parameter values are shown in Appendix 2 Table 1.

**Appendix 2 Figure 1.**
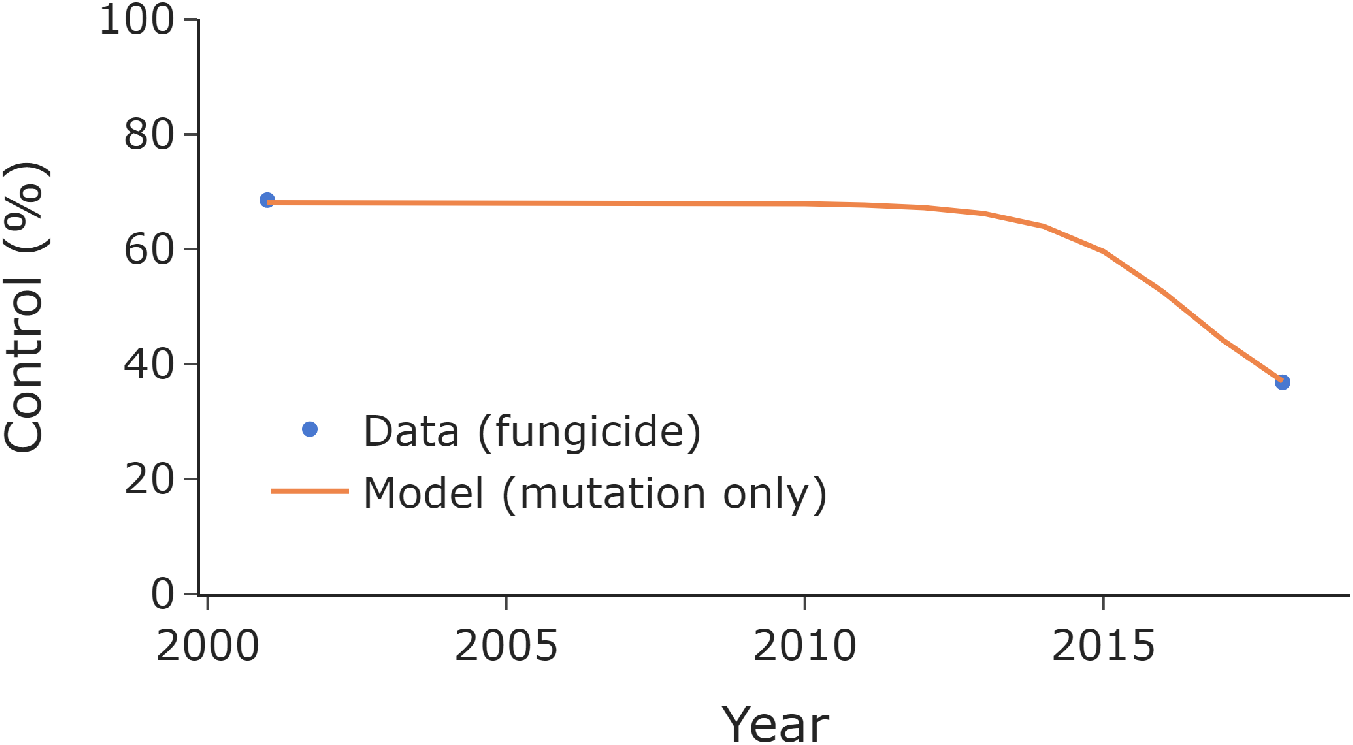
Mutation scale fitting. We found the optimal initial trait value and mutation scale that led to the model matching the level of control achieved in the first and last year. The initial trait value determined the level of control initially, and the mutation scale affected the rate of loss of control. The resulting mutation scale is shown in Appendix 2 Table 1. The value found is an upper bound for the mutation scale, since this is the value in the scenario where there is no initial standing variation in the pathogen population. All other model runs involve initial standing variation and a lower value for the mutation scale (assumed to be 10% of the upper bound), as per the model fit in the main text (Figure 2). Here *n*_*k*_ = 500.

**Appendix 2 Table 1.** Mutation parameters used in the model, reported to 3 significant figures.

| Parameter | Value |
| --- | --- |
| Mutation proportion | $1.23 \times 10^{-5}$ |
| Mutation scale upper bound | 0.0198 |
| Mutation scale used in the model | 0.00198 |

## Appendix 3 Testing the assumption that the mutation scale was 10% **of the previously found mutation scale upper bound**

Here we test the assumption that the mutation scale was 10% of the upper bound mutation scale value found in Appendix 2. We test 4 possible values: 1%, 10%, 50% and 90%. For each value, we fit a fungicide distribution, finding the optimal parameters using the Optuna package (as in the main text).

All values for the proportion of the maximum mutation scale achieve a comparable model fitting score on the training sets (Appendix 3 Figure 1), although the shape of the fits vary slightly in each case (Appendix 3 Figure 2). The model fitting scores on the test set vary slightly (Appendix 3 Figure 1).

**Appendix 3 Figure 1.**
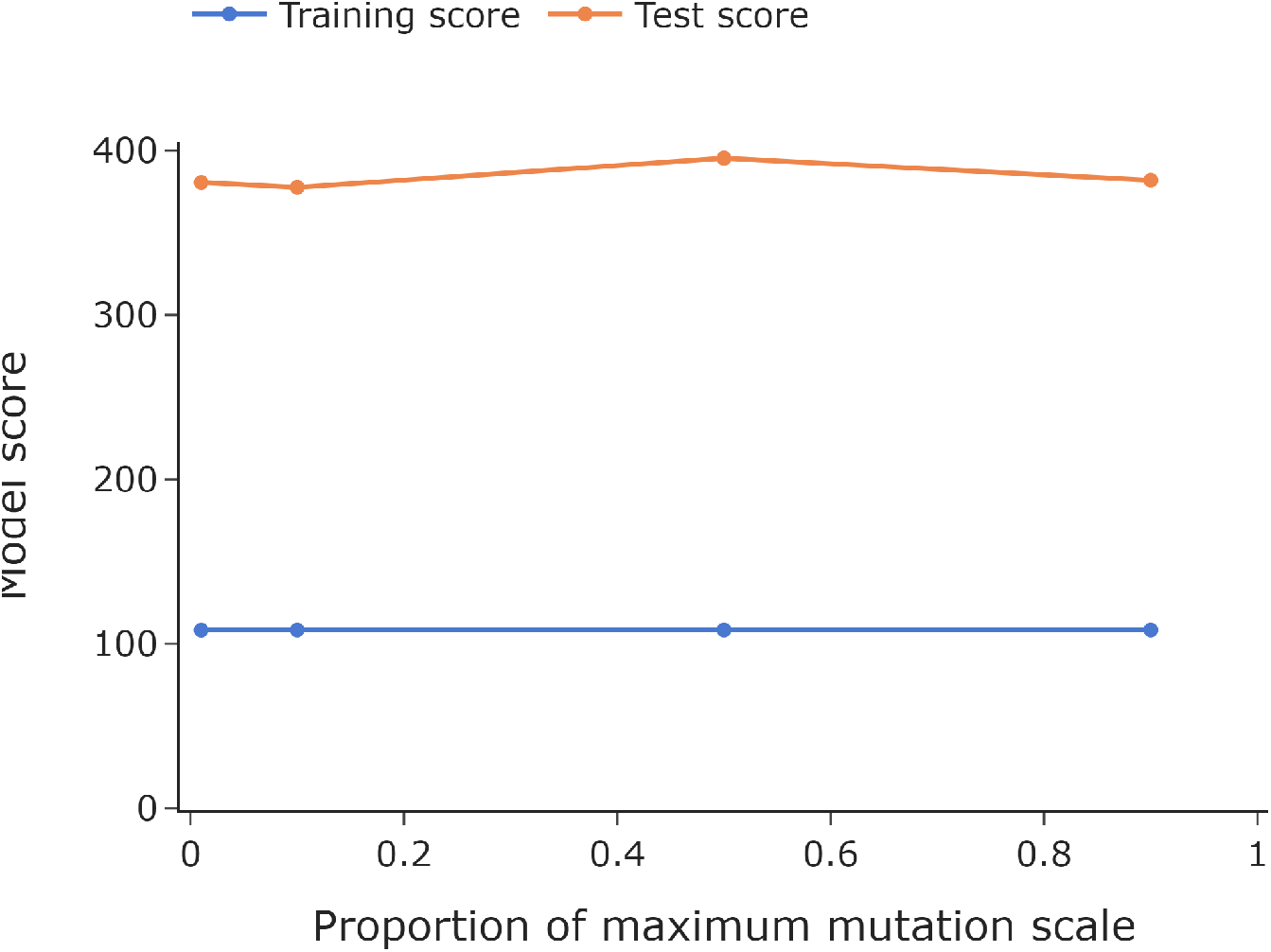
Model fit scores. The model fit score here is the mean of the squared residuals between the model output and the fungicide control data for the optimal fungicide distribution parameters found using Optuna. We show the values on the training set (2001 to 2012) and on the test set (2013 to 2018). Lower values indicate a better fit. The performance on the training set is better than the test set, but all mutation scales lead to similar model performance.

**Appendix 3 Figure 2.**
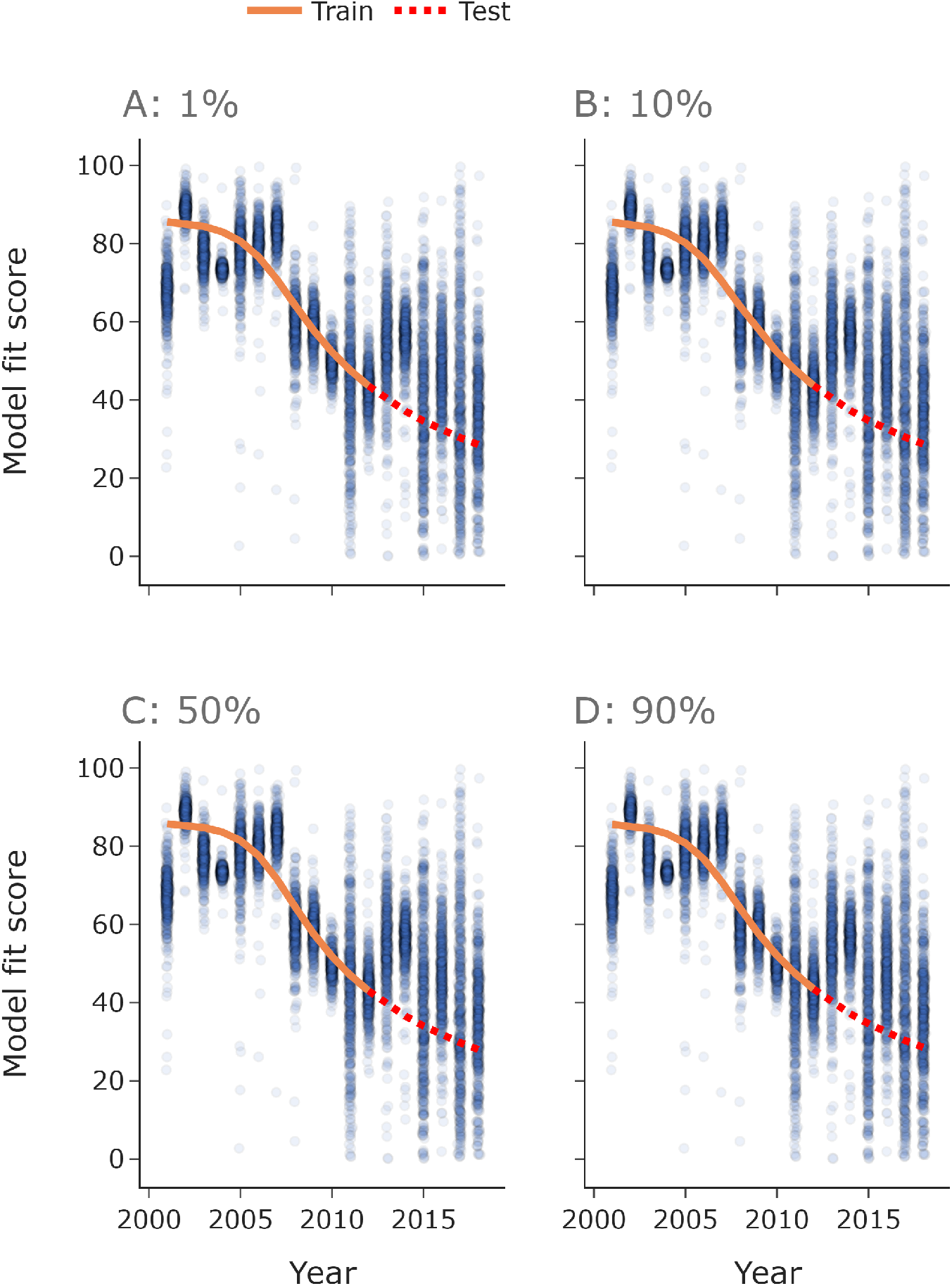
Model fits. For each mutation scale (1%, 10%, 50% and 90% of the mutation upper bound from Appendix 2), we fit the model using Optuna. The optimal model fits are shown here. Here *n*_*k*_ = 500.

The model outputs are qualitatively very similar (Appendix 3 Figure 3). The optimal model parameters are quite similar across the different model fits (Appendix 3 Table 1).

**Appendix 3 Figure 3.**
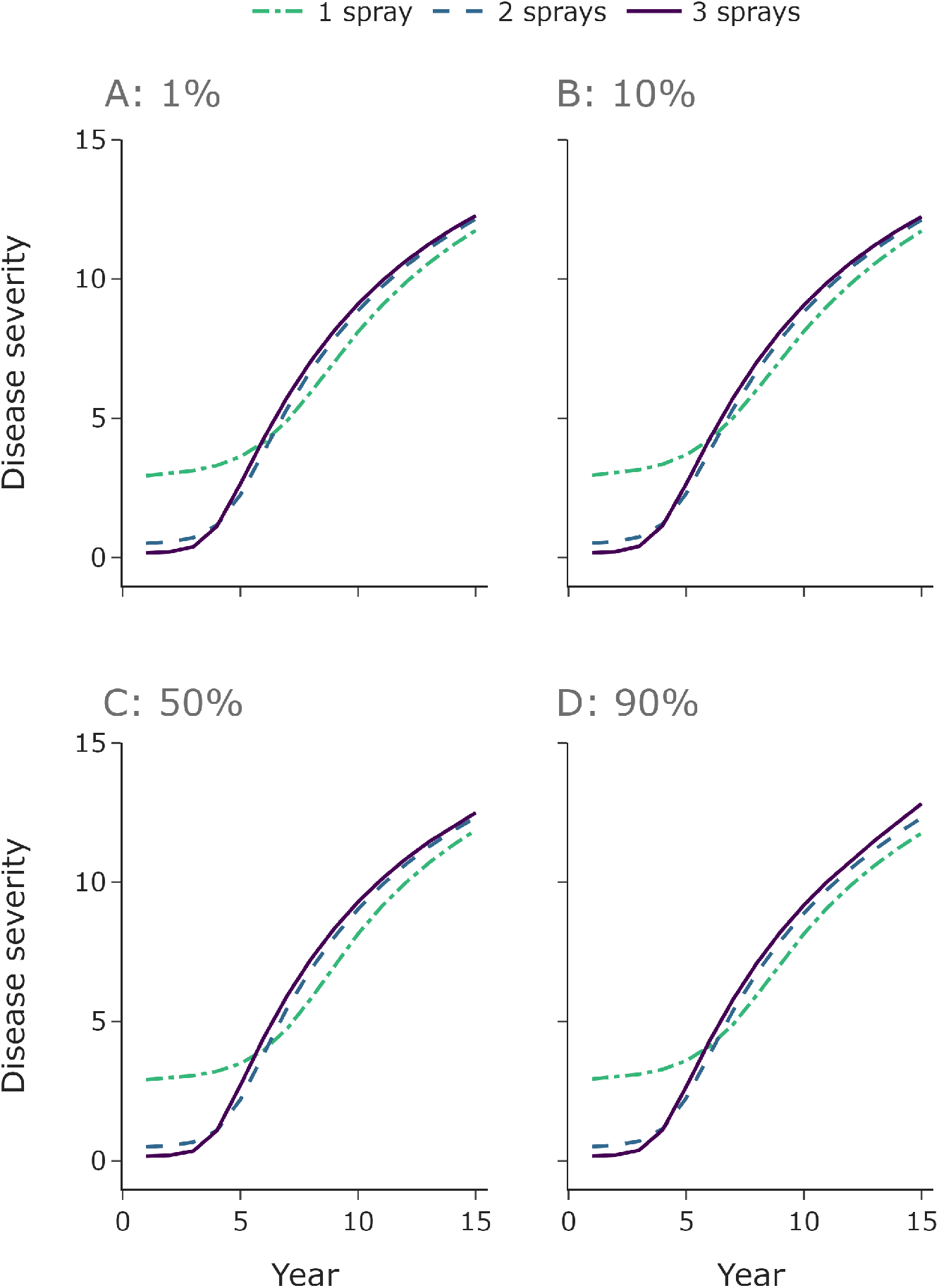
Model outputs. The model outputs for all percentage values are qualitatively similar. Here we use the median value for the infection rate *β*_0_ in every year. The parameters values are in Appendix 3 Table 1. Here *n*_*k*_ = 500.

**Appendix 3 Table 1.** Optimal distribution parameters for each mutation scale, quoted to 3 significant figures.

| % of mutation scale upper bound | Distribution mean | Dist. shape parameter |
| --- | --- | --- |
| 1% | 9.69 | 0.816 |
| 10% | 9.44 | 0.843 |
| 50% | 10.4 | 0.745 |
| 90% | 9.84 | 0.801 |

## Appendix 4 Example distributions – varying the mean but keeping the shape parameter fixed

In Figure 6 (main text) we test how outcomes vary with different cultivar efficacies depending on the initial mean trait value. Appendix 4 Figure 4 shows some examples of the initial host distributions used, alongside the distribution corresponding default host trait mean used in the other model runs.

**Appendix 4 Figure 1.**
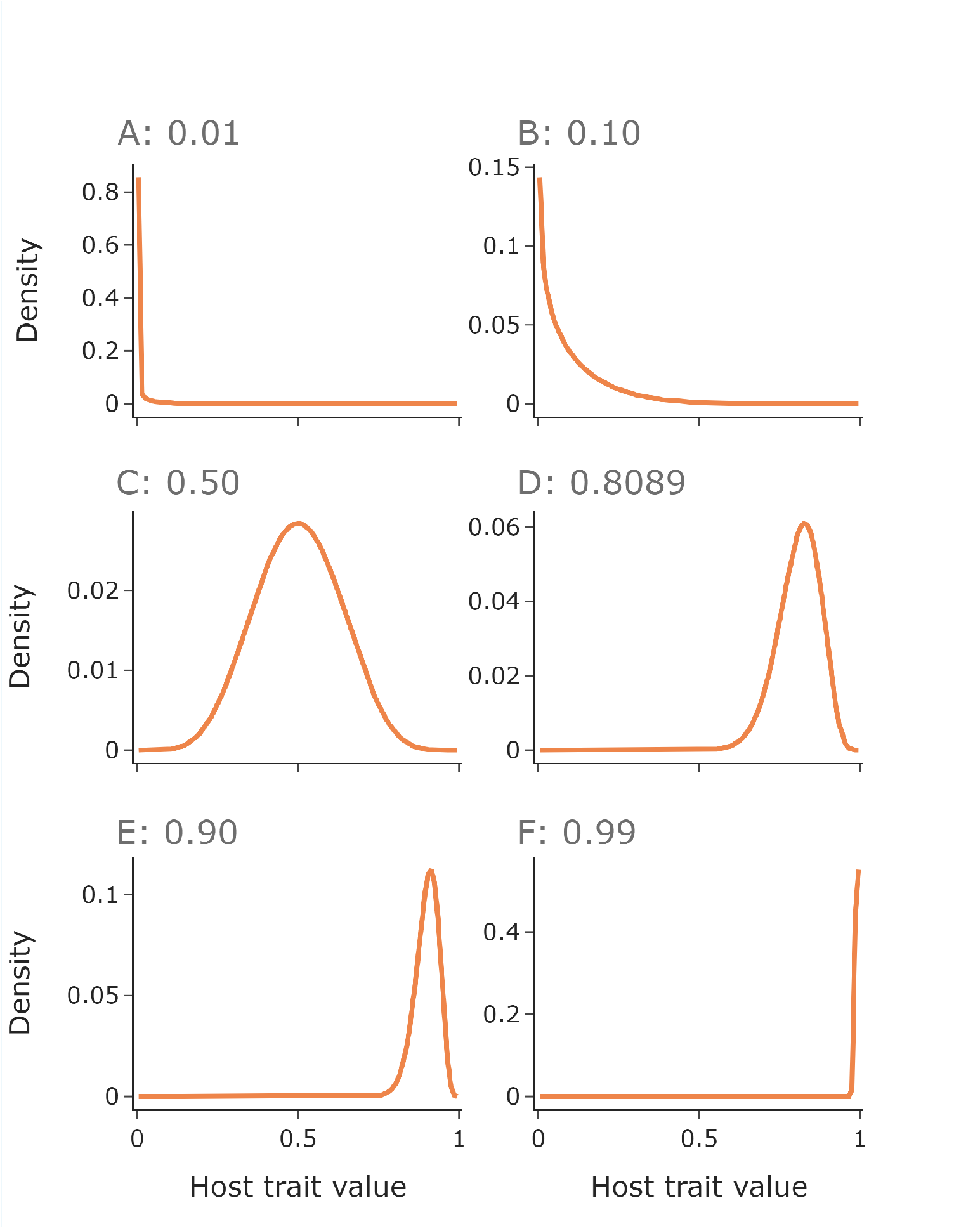
Example initial host distributions used in Figure 6. We show 6 example host distributions. In Figure 6 the initial trait mean varies from 0.01 (shown in **A**) to 0.99 (shown in **A**). For comparison, we show the initial distribution for the default mean host trait value in panel **D**.

